# Ciliary sensing in tanycytes couples nutrient availability to metabolic regulation

**DOI:** 10.64898/2025.12.31.697168

**Authors:** Manon Rivagorda, Constance Kienle, Akila Chandrasekar, Sara Dori, Sofia Grünberg, Sreekala Nampoothiri, Ümit Özorhan, Wiebke Brandt, Jonas Rotter, Nina Feller, Vanessa Neve, Surya Rai, Sonja Binder, Ingo Bechmann, Rubén Nogueiras, Helge Müller-Fielitz, Vincent Prévot, Markus Schwaninger

**Affiliations:** Institute for Experimental and Clinical Pharmacology and Toxicology, Center of Brain, Behavior and Metabolism, University of Lübeck, Lübeck, Germany; Univ. Lille, Inserm, CHU Lille, Laboratory of Development and Plasticity of the Neuroendocrine Brain, Lille Neuroscience & Cognition, UMR-S 1172, DISTALZ, EGID, Lille, France; Institute of Anatomy, Leipzig University, Leipzig, Germany; Department of Physiology, CIMUS, University of Santiago de Compostela-Instituto de Investigación Sanitaria, Santiago de Compostela, Spain

## Abstract

Body homeostasis relies on accurate communication between the brain and the periphery. Disruption of this communication can contribute to disease. Tanycytes are located at the interface of the cerebrospinal fluid (CSF), bloodstream, and hypothalamus, where they sense circulating nutrients and regulate neuroendocrine axes and metabolism. However, the mechanisms by which they sense CSF signals remain largely unclear. Recent evidence that tanycytes possess primary cilia — key sensory organelles — led us to hypothesize that tanycytic cilia function as sensory antennae that detect metabolic cues in the CSF. Here, we demonstrate that tanycytic cilia exhibit distinct morphologies across subtypes and physiological states. They respond dynamically to hormonal and nutrient availability; notably, excess oleic acid shortens cilia, promotes lipid droplet accumulation, and reduces Ca²⁺ responses to ATP and glucose. Disrupting cilia via knockdown of intraflagellar transport (IFT) genes produced similar defects and impaired autophagy. Finally, selective *Ift88* knockout in tanycytes increased body weight and reduced thermogenic activity in female mice. These findings identify tanycytic cilia as key sensors regulating energy balance.

## Introduction

Whole-body physiology relies on finely tuned interactions between the brain and the periphery. The organism’s ability to adapt to a constantly changing environment depends on the brain’s capacity to accurately detect, interpret, and integrate signals from the periphery^1^. Any perturbation in this intricate communication network can lead to maladaptive responses and chronic diseases, such as cognitive decline and metabolic disorders.

One of the key cellular structures enabling neurons and glial cells to detect internal and environmental stimuli is the primary cilium. This solitary organelle acts as a cellular antenna that senses changes in the extracellular milieu^2,3^. Primary cilia are enriched in specific signalling components, notably G protein–coupled receptors (GPCRs), which accumulate at the ciliary membrane and initiate distinct intracellular pathways upon ligand binding^4,5^. Thereby, primary cilia can sense hormones, growth factors, nutrients, or mechanical forces such as fluid flow. The functional integrity of primary cilia is therefore critical for homeostasis, and ciliary dysfunction leads to severe phenotypes including memory impairment, obesity, and diverse metabolic syndromes^6–11^. Indeed, primary cilia in the hippocampus and hypothalamus have been shown to regulate key physiological functions including appetite and energy balance^12–14^.

A particularly important brain region for coordinating brain–body communication is the hypothalamus, especially the area surrounding the third ventricle (3V) and the median eminence (ME). In this region, specialized ependymo-glial cells known as tanycytes line the walls and floor of the 3V. While their cell bodies are located in the ventricle wall and contact the cerebrospinal fluid (CSF), they send long processes into the parenchyma. Based on the target of their projections, tanycytes are classified into α1-(dorsomedial and ventromedial hypothalamic nuclei), α2-(dorsomedial arcuate nucleus (ARH)), β1-(ventromedial ARH), and β2-tanycytes (fenestrated vessels in the ME). The privileged position of tanycytes between bloodstream, CSF and mediobasal hypothalamus (MBH) enables numerous interactions with the centers that regulate neuroendocrine axes and metabolism. Thus, tanycytes control the function of proopiomelanocortin (POMC) and neuropeptide Y (NPY) neurons in the ARH^15–17^. Moreover, they transport metabolic hormones, such as insulin, leptin, FGF21, GLP-1, and ghrelin, across the blood-brain barrier to their central site of action^17–22^. Finally, tanycytes play crucial roles in regulating neuroendocrine axes. They control the access of neurosecretory nerve terminals to the portal pituitary vessels in the ME^23–25^. To accomplish these functions, tanycytes must sense hormonal and metabolic cues from the circulation and the CSF. While much is known about how tanycytes detect blood-borne signals through their basal processes that contact fenestrated capillaries, it remains unclear how these cells sense and respond to cues presented by the CSF to their apical surface.

A strong candidate for this sensory function is the primary cilium. Recent studies^26,27^ have described the presence of primary cilia on tanycytes, challenging the traditional view that these cells are non-ciliated, in contrast to neighbouring ependymal cells bearing multiple motile cilia that promote CSF flow. Bi- and uni-ciliated ependymal cell types lining the third and fourth ventricles were shown to correspond to specific subtypes of tanycytes originating from anterior ventral neural progenitors^27^. Newborn Siberian hamster pups exposed in utero and postnatally to long photoperiods exhibited increased numbers of ciliated tanycytes in the hypothalamic ependymal region, implicating these cilia-bearing cells in seasonal growth and reproductive programming^26^. However, the regulation of cilia morphology and their physiological functions in tanycytes are still unknown.

Given the well-established role of primary cilia as sensors of environmental changes, we hypothesized that tanycytic cilia may serve as antennae that detect nutrient and hormonal fluctuations within the CSF, enabling adaptive cellular responses and facilitating the regulation of metabolism.

Here, we demonstrate that tanycytes possess cilia with distinct morphological characteristics that vary among their different subtypes. The morphology of tanycytic cilia also varies depending on the physiological context, including sex and age. Furthermore, we discovered that tanycytic cilia are dynamically regulated by hormonal and nutrient availability. Specifically, cilia length decreases in the presence of excess oleic acid, coinciding with lipid droplet accumulation in tanycyte cell bodies and a reduction in ATP-evoked Ca²⁺ signalling. Using an in vitro model of ciliary impairment, we observed comparable blunting of Ca²⁺ responses to both ATP and glucose, confirming that cilia are required for sensing these nutrients. Moreover, tanycytic cilia impairment in vitro decreased the autophagic flux, likewise resulting in lipid droplet accumulation in tanycytes. Finally, we demonstrate that impairing cilia selectively in adult tanycytes leads to increased body weight, decreased core body temperature, and reduced brown adipose tissue (BAT) activity specifically in female mice.

## Results

### Human and mouse tanycytes are ciliated with subtype-specific morphologies

Consistent with one previous study^27^, we found that mouse tanycytes extend cilia from their cell bodies toward the ventricular lumen, where they directly contact the CSF (**Fig. 1a**). Also, human tanycytes carry cilia on their ventricular surface (**Fig. 1c**). A comparative analysis of single-cell RNA sequencing of tanycytes from mice and humans revealed subcluster-specific profiles of ciliary gene expression (**Fig. 1b,d, and Extended Data Fig. 1a**)^28^. In both the species, subcluster T3 (mT.3 and hT.3) that correspond to α1-tanycyte subtype were enriched for key ciliary genes including ADP-ribosylation factor-like GTPase 13B (*Arl13b* / *ARL13B)* while clusters 6-8 (mT.6-8, hT.6-8) that correspond to β-tanycytes exhibited reduced expression of *Arl13b* / *ARL13B*, suggesting that the molecular composition of tanycytic cilia varies according to their position within the 3V.

**Fig. 1.**
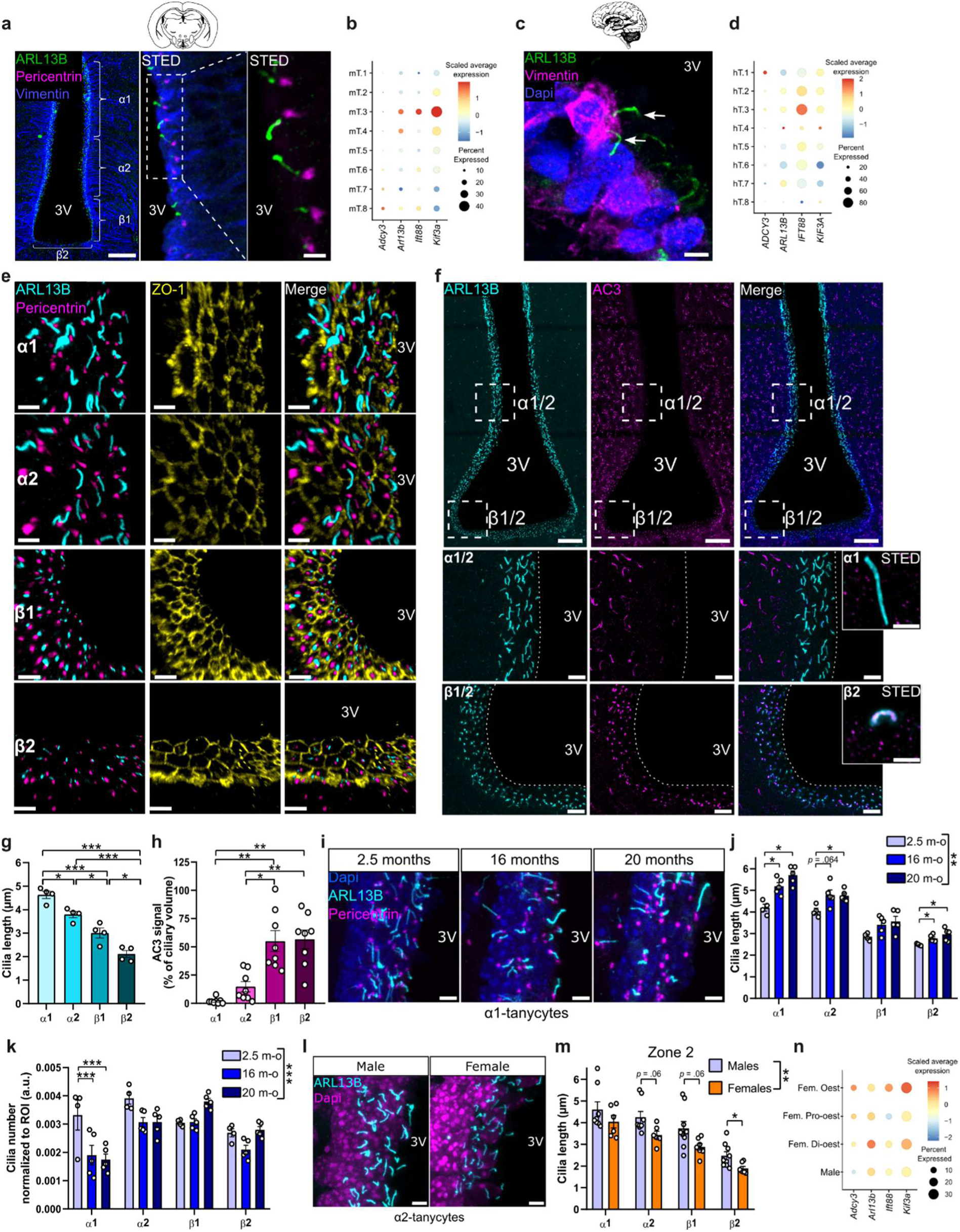
Tanycytes bear cilia with distinct morphologies across subtypes and physiological states. **a,** Immunofluorescent staining of tanycytes (vimentin, blue), cilia (ARL13B, green) and basal bodies (pericentrin, magenta) in a coronal section of a mouse brain. Scale bar, 100 µm. Tanycyte subtypes are indicated in white. The zoomed-in images were generated by STED microscopy and show the extension of cilia from tanycyte cell bodies to the third ventricle (3V). Scale bar, 2 µm. **b,** Single-cell RNA sequencing data of *Adcy3, Arl13b, Ift88* and *Kif3a* in the tanycyte subgroups in mice. **c,** Immunofluorescent staining of tanycytes (vimentin, magenta) and cilia (ARL13B, green) in a coronal section of a human brain (representative image). Arrows indicate tanycytic cilia. Scale bar, 5 µm. **d,** Single-cell RNA sequencing data of *ADCY3*, *ARL13B*, *IFT88* and *KIF3A* in the tanycyte subgroups in humans. **e,** Immunofluorescent stainings of cilia (ARL13B, cyan), basal body (pericentrin, magenta) and tight junctions (ZO-1, yellow) in a coronal section of a mouse brain, representative images of each tanycytes subtypes (α1, α2, β1 and β2). Scale bar, 5 µm. **f,** Immunofluorescent stainings of cilia (ARL13B, cyan) and “neuronal” cilia marker (AC3, magenta) in a coronal section of a mouse brain. Upper panel shows the overview of the 3V (scale bar, 100 µm) and the dashed-line squares indicate the zoomed images in the α1/α2 and β1/β2 transition zones, which are shown in the bottom panels (scale bar, 10 µm). The dotted line indicates the tanycyte-3V border. Bottom right panel shows representative α1- and β2-tanycytes with STED imaging (scale bar, 2 µm). **g,** Cilia length measurement in the 4 tanycyte subtypes shown in (e). The length of the cilia was measured from control mice and quantified in 3D using the Imaris software. **h,** Quantification of AC3 signals within tanycytic cilia (ratio of AC3 volume to ARL13B volume, expressed as percentage), in the 4 tanycyte subtypes from control mice (related to f). **i,** Cilia and basal body immunofluorescent staining (ARL13B, cyan; pericentrin, magenta) in α1-tanycytes of young (2.5 months) and aged (16 and 20 months) male control mice. Scale bar, 5 µm. **j,** Cilia length and **k** cilia number normalized to ROI in the 4 tanycyte subtypes from young and aged adult control mice. **l,** Cilia immunofluorescent staining (ARL13B, cyan) in α2-tanycytes of male and female control mice. Scale bar, 5 µm. **m,** Cilia length in the 4 tanycyte subtypes in male and female control mice, in the tanycyte zone 2 of the anteroposterior axis (in the ME). **n,** Single-cell RNA sequencing data of *Adcy3, Arl13b, Ift88* and *Kif3a* in the tanycyte subgroups in male and female oestrus, pro-oestrus and dioestrus mice. All data are expressed as mean ± s.e.m., ∗, *p* < 0.05; ∗∗, *p* < 0.01; ∗∗∗, *p* < 0.001. *n* denotes the number of mice. Detailed information on the test statistics is provided in **Extended Data Table 4**.

Since ARL13B is a key structural protein of the ciliary axoneme, we tested the notion that cilia morphology varies between tanycyte subpopulations by staining ARL13B, basal bodies (pericentrin), and the tight junction protein zonula occludens-1 (ZO-1) along the dorso-ventral axis of the mouse 3V. ARL13B delineates the contour of cilia and ZO-1 the apical cell borders. Alpha1-tanycytes had the longest cilia (mean ± SEM, 4.6 ± 0.2 µm), while β2-tanycytes had the shortest (mean ± SEM, 2.1 ± 0.2 µm) (**Fig. 1e,g**). The proportion of ciliated tanycytes remained constant in all subtypes (**Extended Data Fig. 1c**). To evaluate cilia length along the antero-posterior axis, we divided the 3V into four zones according to the ventricular shape and the surrounding hypothalamic structures (**Extended Data Fig. 1d**)^29^. The subtype-specific distribution of cilia length was preserved throughout the zones (**Fig. 1m and Extended Data Fig. 1e-g**).

While we occasionally observed tanycytes containing two basal bodies, two cilia were rarely present, if at all (<10%, data not shown); the vast majority of α-tanycytes remained uni-ciliated in contrast to the multiple cilia in ependymocytes (**Fig. 1e and Extended Data Fig. 1b**). Progressing toward the β-region, the characteristic honeycomb pattern revealed by ZO-1 staining enabled unambiguous assignment of β-tanycytes, which were uniformly uni-ciliated, each bearing a single basal body and a short primary cilium projecting into the ventricle (**Fig. 1e**).

Our single-cell data analysis also showed enriched expression of adenylate cyclase 3 (*Adcy3*) in mouse β-tanycytes (**Fig. 1b**). AC3, a cAMP-generating enzyme known to be highly concentrated in neuronal cilia^30^, displayed a similar pattern at the protein level. While AC3 was uniformly expressed in neuronal cilia throughout the parenchyma, its expression in tanycytic cilia was strongest in β-tanycytes and progressively diminished toward the dorsal 3V (**Fig. 1f,h**). These findings support the idea that the signalling compartments formed by tanycytic cilia are tailored to the specific functional identity of each tanycyte subtype. The pronounced enrichment of AC3 in β1- and β2-tanycytes suggests active cAMP signalling within the floor of the 3V, adjacent to the ME and ARH, whereas AC3 signalling pathways appear less prominent in the upper ventricular regions. Whether this pattern is conserved in humans is unclear, because the transcript levels approached the limit of detection in the snRNAseq study **(Fig. 1d)**.

### Tanycytic cilia morphology varies by physiological context including age and sex

Consistent with previous findings on morphological abnormalities of neuronal cilia in aging^11,31^, we observed longer tanycytic cilia in 16- and 20-month-old mice than in young adults (2- to 3-month-old) (**Fig. 1i,j**). This increase in cilia length is observed in all tanycyte subtypes, even though the subtypes maintained their specific length differences in old age. Interestingly, the elongation was accompanied by a reduction in the number of ciliated tanycytes, specifically in the α1 region (**Fig. 1i,k**).

Interestingly, we observed consistently shorter cilia in female compared to male mice across all tanycyte subtypes (**Fig. 1l,m**). This sexual dimorphism in cilia morphology was restricted to the zones 2 and 3 of the antero-posterior axis, corresponding to the ME, where tanycytes are positioned to sense circulating sex hormones (**Extended Data Fig. 1e-k**)^16^. Since ME tanycytes have been identified as key integrators of peripheral sex hormone signals to the hypothalamus, and since oestrogens are known to modulate primary cilia length in other tissues, our results raise the possibility that tanycytic cilia may be modulated by peripheral oestrogen signalling^16,32^. In support of this hypothesis, we found longer tanycytic cilia during the proestrus, when oestrogen levels are at their highest, and oestrous stages of the ovarian cycle^33^, than during dioestrus, when progesterone concentrations increase (**Extended Data Fig. 1l-n**). Our single-cell data analysis showed that *Adcy3* is expressed at lower levels in male tanycytes, and that *Ift88* as well as other ciliary genes are highly sensitive to oestrogens, with their expression being down-regulated during proestrus. This corresponds to the sexual dimorphism of ciliary morphology (**Fig. 1n**). Together, these morphological and molecular differences among tanycyte subtypes suggest functional specializations of tanycytic cilia that are regulated by the physiological context.

### Morphology of tanycytic cilia is modulated by nutrient and hormonal status

#### TSH and thyroid hormones

Tanycytes express thyroid hormone (TH) transporters, receptors, and metabolizing enzymes, consistent with the notion that they mediate metabolic effects of thyrotropin (TSH) and TH^34^. Since a recent study in Siberian hamsters suggested a link between elevated triiodothyronine (T3) production in the MBH during long photoperiods and longer tanycytic cilia^26,35^, we tested whether thyroid hormones directly modulate the length of tanycytic cilia that were visualized by staining axonemes (ARL13B) and basal bodies (pericentrin). In primary tanycytes, T3 treatment for 16 hours showed a non-significant trend toward longer cilia and L-thyroxine (T4) and TSH increased cilia length in comparison to hypothyroid conditions (**Fig. 2a,b**). The proportion of cells bearing cilia remained unchanged (**Fig. 2a,c**). These results suggest that the length of tanycytic cilia reflects the supply of thyroid hormones that control metabolism, potentially by modulating ciliary signalling.

**Fig. 2.**
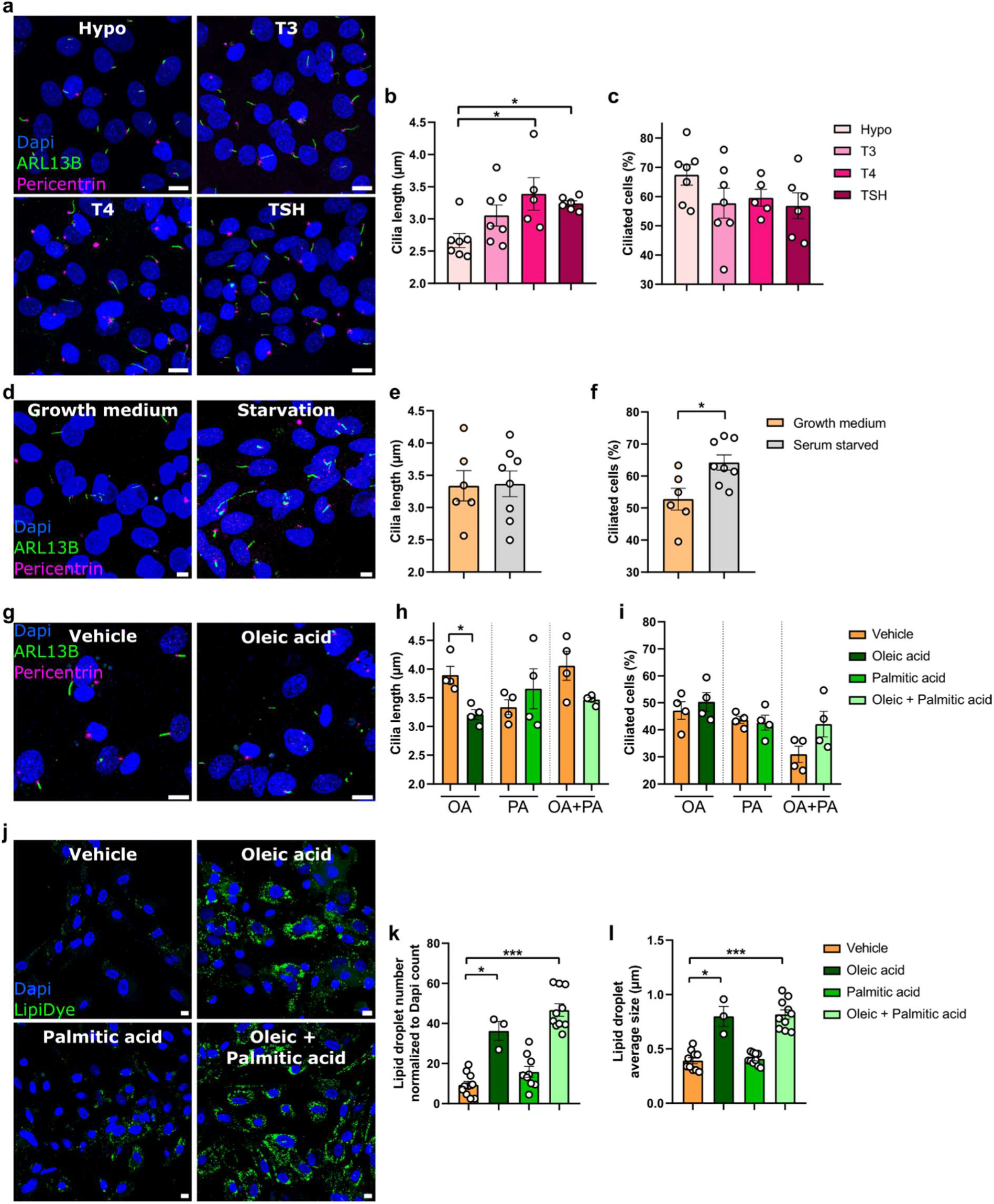
Metabolic and endocrine cues remodel tanycytic cilia morphology. **a,** Cilia and basal body immunofluorescent staining (ARL13B, green; pericentrin, magenta) in primary tanycytes treated with thyroid hormone-depleted medium (= Hypo), T3 (10 nM), T4 (10 nM) or TSH (60 mIU) for 16 h. Scale bar, 10 µm. **b,c,** Cilia length **(b)** and percentage of ciliated tanycytes **(c)** in primary tanycytes treated with thyroid hormone-depleted medium (= Hypo), T3 (10 nM), T4 (10 nM) or TSH (60 mIU) for 16 h. **d,** Cilia and basal body immunofluorescent staining (ARL13B, green; pericentrin, magenta) in primary tanycytes treated with normal growth medium or starvation medium for 16 h. Scale bar, 5 µm. **e,f,** Cilia length **(e)** and percentage of ciliated tanycytes **(f)** in primary tanycytes treated with normal growth medium or starvation medium for 16 h. **g,** Cilia and basal body immunofluorescent staining (ARL13B, green; pericentrin, magenta) in primary tanycytes treated with vehicle or oleic acid (OA, 100 nM) for 6 h. Scale bar, 10 µm. **h,i,** Cilia length **(h)** and percentage of ciliated tanycytes **(i)** in primary tanycytes treated with vehicle, OA (100 µM), palmitic acid (PA, 100 µM), or a combination of OA and PA for 6 h. **j,** Lipid droplet dye (Lipidye, green) in primary tanycytes treated with vehicle, OA (100 µM), PA (100 µM), or a combination of OA and PA for 6 h. Scale bar, 10 µm. **k,l,** Lipid droplet number per cell (per Dapi count) **(k)** and size **(l)** in primary tanycytes treated with vehicle, OA (100 µM), PA (100 µM), or a combination of OA and PA for 6 h. All data are expressed as mean ± s.e.m., ∗, *p* < 0.05; ∗∗, *p* < 0.01; ∗∗∗, *p* < 0.001. *n* denotes the number of wells. Detailed information on the test statistics is provided in **Extended Data Table 4**.

#### Starvation and excess fatty acid

Primary cilia are equipped with a diverse repertoire of receptors and function as sensors of metabolic cues^13,36–38^. Thereby, they convey information about the organism’s nutritional state^36,39,40^. Since tanycytes form the metabolic gateway to the hypothalamus and carry primary cilia, we asked whether tanycytic cilia respond to changes in nutrient availability. To approach this question, we examined whether ciliary morphology, in primary tanycytes, is affected by serum deprivation. Starvation medium (without serum) resulted in an increase in ciliated tanycytes (**Fig. 2d,f**), while cilia length remained unchanged (**Fig. 2d,e**). These findings indicate that primary tanycytes respond to the lack of serum by promoting ciliogenesis, consistent with observations in other cell types where serum and nutrient deprivation enhances cilia formation^41,42^.

Since tanycytes act as gatekeepers of hypothalamic lipid metabolism^43,44^, we wondered whether the serum effect could be at least partly due to its fatty acid content. We found that primary tanycytes exposed to the monounsaturated fatty acid oleic acid, for 6 hours, exhibited significantly shorter cilia without changes in the proportion of ciliated cells (**Fig. 2g-i**). In contrast, the saturated fatty acid palmitic acid had no effect on tanycytic cilia morphology and number. Oleic acid treatment, alone or with palmitic acid, markedly increased both the number and size of lipid droplets, the storage organelles of neutral lipids, whereas palmitic acid alone had no effect (**Fig. 2j-l**). These findings align with a different distribution of oleic acid and palmitic acid in tanycytes and with in-vivo data showing that high-fat diets induce lipid droplet accumulation in tanycytes^44,45^. Whereas starvation increases the proportion of ciliated tanycytes, oleic acid (an unsaturated fatty acid) triggers cilia shortening, consistent with a shift in ciliary signalling under conditions of metabolic surplus. Palmitic acid (a saturated fatty acid) showed no effect, indicating that ciliary responses may distinguish between unsaturated and saturated lipid loads.

### Tanycytic cilia are important mediators of the downstream signalling pathways underlying metabolic control

Tanycytes are known to be important sensors of blood glucose levels. Various mechanisms have been identified^15,46–49^, one of which is that tanycytes indirectly sense glucose by responding to the glucose-derived ATP. This propagates the glucose response through Ca^2+^ waves within the tanycytic layer. We investigated whether fatty acids could impair the Ca^2+^ response to ATP. Indeed, the increase in intracellular Ca^2+^ levels in response to ATP stimulation was significantly reduced after 6 hours of exposure to oleic acid alone or combined with palmitic acid (**Fig. 3a-c**). Because palmitic acid has no additive effect, this demonstrates that an excess of oleic acid, as observed in metabolic syndrome, interferes with tanycyte sensitivity to ATP, possibly resulting in less efficient signalling than in physiological conditions.

**Fig. 3.**
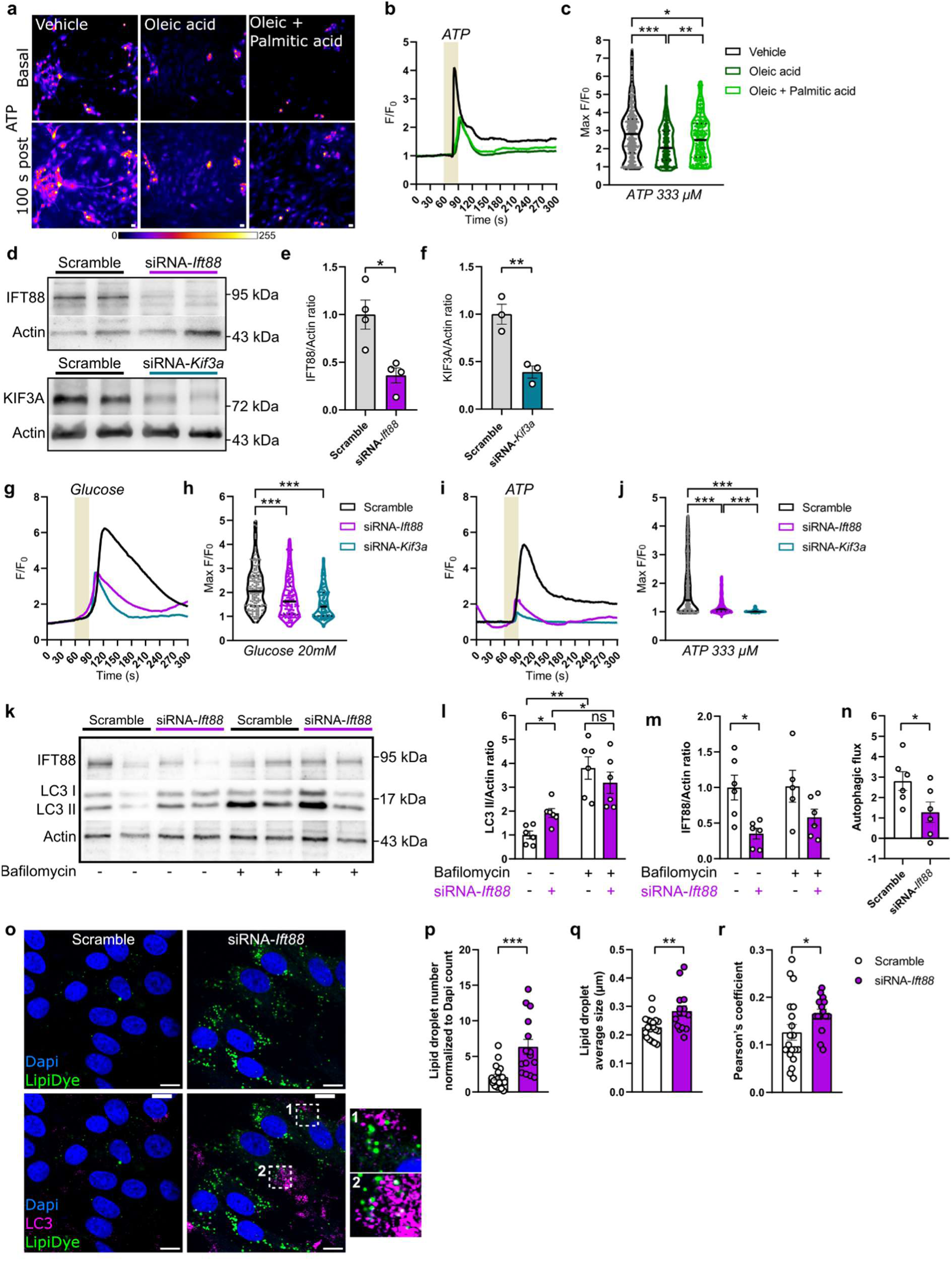
Ciliary disruption impairs Ca²⁺ signalling and autophagic flux, leading to an accumulation of lipid droplets in tanycytes. **a,** To measure intracellular calcium concentrations ([Ca^2+^]_i_) in primary tanycytes, cells were loaded with the calcium indicator Fluo4. The fluorescence emission at 488 nm, shown in pseudocolours, increased in tanycytes after stimulation with ATP (333 µM). Scale bar, 100 µm. **b,** Representative trace of the fluorescence ratio (F) relative to the baseline ratio (F_0_) after stimulation with ATP (30 s stimulus, beige zone) in primary tanycytes treated with vehicle, oleic acid (OA, 100 µM), or a combination of OA and palmitic acid (PA, 100 µM) for 6 h. **c,** Quantification of ATP-induced [Ca^2+^]_i_ as determined by maximal F/F_0_. **d,** Western blot of primary tanycytes 72 h after transfection with siRNA-scramble, siRNA-*Ift88* or siRNA-*Kif3a*, and **e** quantification of IFT88 and **f** KIF3A, relative to the scramble-treated cells. **g,** Representative trace of F/F_0_ after stimulation with glucose (20 mM, 30 s stimulus, beige zone) in primary tanycytes 72 h after transfection with siRNA-scramble, siRNA-*Ift88* or siRNA-*Kif3a*. **h,** Quantification of glucose-induced [Ca^2+^]_i_ as determined by maximal F/F_0_. **i,** Representative trace of F/F_0_ after stimulation with ATP (333 µM, 30 s stimulus, beige zone) in primary tanycytes 72 h after transfection with siRNA-scramble, siRNA-*Ift88* or siRNA-*Kif3a*. **j,** Quantification of ATP-induced [Ca^2+^]_i_ as determined by maximal F/F_0_. **k,** Western blot of primary tanycytes 72 h after transfection with siRNA-scramble or siRNA-*Ift88*, treated with DMSO of bafilomycin (BafA1, 100 nM) for 4 h, and **l** quantification of LC3 II and **m** IFT88, relative to the vehicle-treated control. **n,** Autophagic flux calculated as the difference in LC3 II levels in presence and absence of BafA1 (100 nM) between Scramble and siRNA-*Ift88* transfected tanycytes. **o,** Lipid droplet and autophagosome staining (Lipidye, green; LC3, magenta) in primary tanycytes 72 h after transfection with siRNA-scramble or siRNA-*Ift88*. Scale bar, 10 µm. **p,q,** Lipid droplet number per cell (per Dapi count) **(p)** and size **(q)** in primary tanycytes 72 h after transfection with siRNA-scramble or siRNA-*Ift88*. **r,** Relative LC3-lipid droplet co-localization was quantified by Pearson’s coefficient analysis. Data are expressed as mean ± s.e.m. (**e,f,l-n,p-r**) or as median and quartiles (**c,h,j**), ∗, *p* < 0.05; ∗∗, *p* < 0.01; ∗∗∗, *p* < 0.001; ns, *p* > 0.05. *n* denotes the number of cells **(c,h,j)**, wells **(e,f,l-n)** or single images analysed from 3 wells **(p-r)**. Detailed information on the test statistics is provided in **Extended Data Table 4**.

Given that excess oleic acid attenuated the Ca^2+^ response in tanycytes and simultaneously disrupted ciliary morphology, and considering that primary cilia are well-established compartments for Ca^2+^ signalling, we hypothesized that tanycytic cilia might be directly required for the Ca^2+^ response to glucose. To test this, we compared intracellular Ca^2+^ dynamics in primary tanycytes with intact cilia to those with impaired cilia. Ciliary disruption was achieved through siRNA-mediated knockdown of *Kif3a* or *Ift88*, two essential components of the anterograde IFT machinery which are essential for cilia function^50,51^. The knockdown of both genes was efficient, as shown at the mRNA and protein level (**Fig. 3d-f and Extended Data Fig. 2a,b**), providing a reliable in vitro model of impaired ciliary signalling.

Remarkably, disruption of tanycytic cilia significantly attenuated the Ca^2+^ responses to both glucose and ATP (**Fig. 3g-j**), demonstrating that tanycytic cilia contribute directly to generating the intracellular signalling in response to glucose and ATP. These findings also support the idea that the defective Ca^2+^ response observed under oleic acid excess may be a consequence of oleic acid-induced ciliary dysfunction.

Several ciliary receptors involved in hypothalamic energy homeostasis signal through cAMP pathways^37^. As AC3 is enriched in tanycytic cilia and catalyses the production of cAMP, these findings suggest that tanycytic cilia are involved in cAMP signalling in β-tanycytes. To test this, we examined cAMP production in primary tanycytes with intact or impaired cilia. Cells were stimulated with forskolin (25 µM), a potent activator of adenylate cyclases, for 10 minutes, after which intracellular cAMP levels were measured. Tanycytes with impaired cilia exhibited a reduced cAMP response compared to control cells (**Extended Data Fig. 2c**). Taken together, these findings demonstrate that tanycytic cilia are essential for proper Ca^2+^ and cAMP signalling in tanycytes.

### Tanycytic cilia are critical for maintaining autophagic flux and lipid balance

Autophagy and cilia frequently interact with each other, albeit in a cell-type-specific manner^11,52–56^. Moreover, tanycytic autophagy-regulated lipid metabolism affects systemic energy balance^45^. Thus, we asked whether impaired tanycytic cilia could disturb the autophagic machinery. We found that LC3 and p62, two key proteins in the autophagy process, accumulated following cilia impairment (**Fig. 3k-n and Extended Data Fig. 2d-f**). Their levels did not further increase with bafilomycin treatment, a lysosomal blocker, indicating a blockage of autophagic flux in tanycytes lacking functional cilia. This suggests that defective autophagic machinery may contribute to the observed metabolic changes.

Lipid droplets are mobilized by a variant of autophagy called lipophagy. To assess whether ciliary dysfunction could mimic the effects of a lipid-rich environment by impairing autophagy, we examined the accumulation of lipid droplets. Knockdown of *Ift88* significantly increased both the number and size of lipid droplets (**Fig. 3o-q**), with increased colocalization of lipid droplets within LC3 puncta (**Fig. 3o,r**), which indicates that intact ciliary signalling is required for correct mobilization of lipid droplets in tanycytes, probably via lipophagy. Altogether, disruption of ciliary function impairs glucose- and ATP-evoked Ca^2+^ responses, alters cAMP dynamics, alleviates the autophagic flux and leads to lipid droplet accumulation. These results identify tanycytic cilia as key regulators of the intracellular signalling pathways that enable tanycytes to sense and respond to metabolic cues.

### Tanycytic cilia impairment increases body weight and reduces core body temperature and brown adipose tissue activity

Since ciliary disruption attenuated Ca^2+^ and cAMP signalling and induced metabolic alterations in primary tanycytes, we assessed whether impairing tanycytic cilia affects systemic metabolic regulation in vivo. To selectively disrupt tanycytic cilia in adulthood, we injected an adeno-associated virus serotype 1/2 (AAV) expressing Cre recombinase and GFP under the tanycyte-specific *Dio2* promoter into the lateral ventricle of *Ift88^fl/fl^* mice^24,57,58^. Two weeks after viral injection, the GFP reporter signal confirmed the efficient and selective targeting of tanycytes (**Fig. 4a**). Immunostaining revealed a significant reduction in IFT88 within the tanycytic layer **(Extended Data Fig. 3a,d**). Quantification of IFT88 specifically inside cilia showed a ∼30% decrease in IFT88 puncta, without changes in cilia length or number (**Fig. 4b,c and Extended Data Fig. 3a-c,e,f**), consistent with a functional impairment of the intraflagellar transport, thus impairing cilia activity, rather than a loss of cilia.

**Fig. 4.**
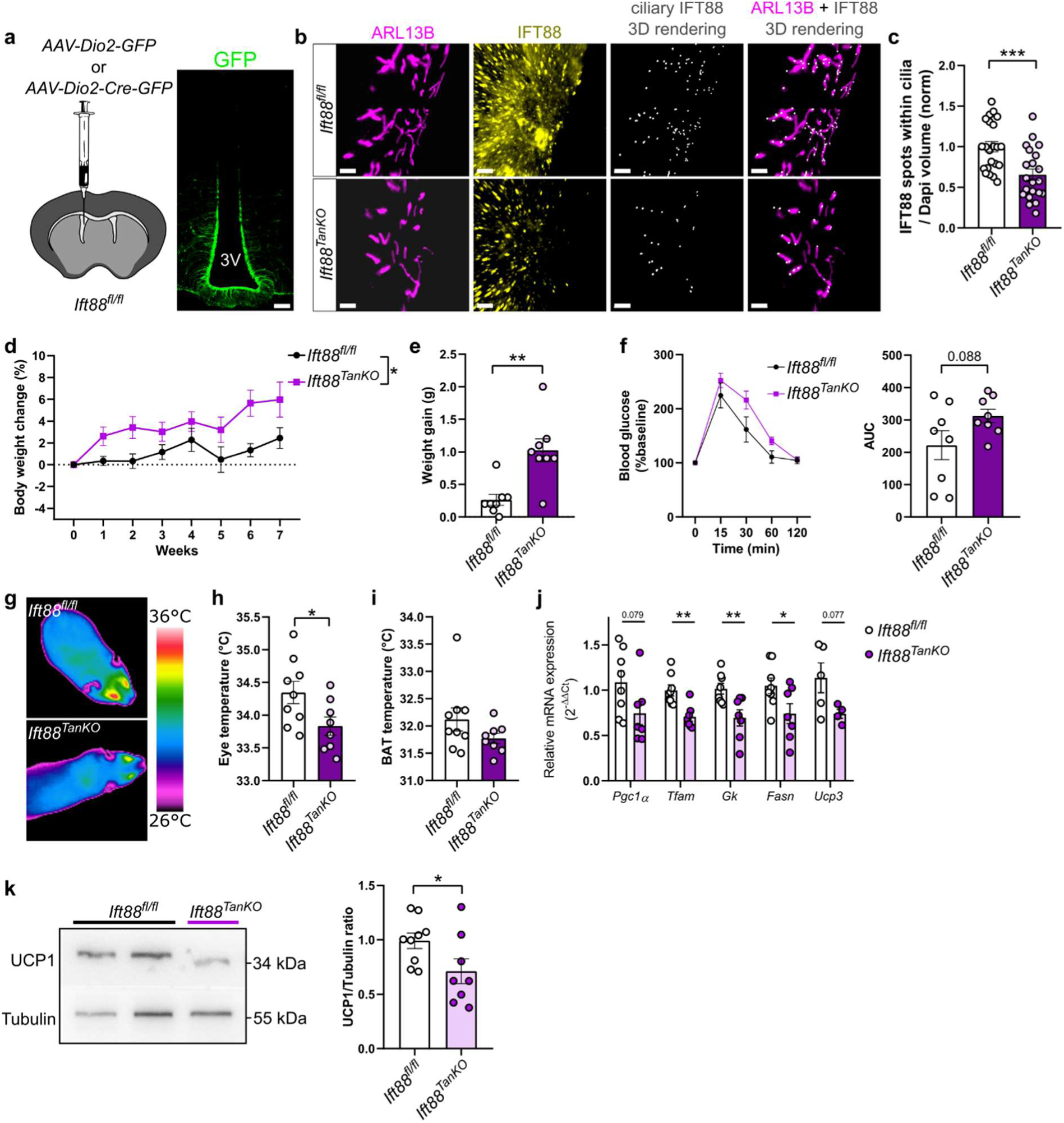
Impairment of tanycytic cilia in female mice results in weight gain accompanied by reduced core body temperature and brown adipose tissue (BAT) activity. **a,**Scheme representing the mouse model *Ift88^fl/fl^* in which we injected the *AAV-Dio2-GFP* for controls or *AAV-Dio2-Cre-GFP* to impair ciliary function in adult tanycytes. Representative fluorescent image of the 3V in a coronal section 3 weeks after injections with *AAV-Dio2-GFP* in the lateral ventricle of *Ift88^fl/fl^* adult mice, showing the tanycytic expression of the virus. Scale bar, 100 µm. **b,** Representative images of tanycytic cilia (ARL13B, magenta) and IFT88 (yellow) stainings as well as ciliary IFT88 3D rendering (grey) in coronal sections of the 3V, after injection of *AAV-Dio2-GFP* or *AAV-Dio2-Cre-GFP* in the lateral ventricle of *Ift88^fl/fl^* adult mice. Scale bar, 5 µm. **c,** Quantification of IFT88 spots within cilia per ROI (versus *AAV-Dio2-GFP*) in both males and females. **d,** Body weight change (%) from the week of injection to the week 7 after induction of the tanycytic cilia impairment in female mice. **e,** Weight gain 7 weeks after induction of the tanycytic cilia impairment in female mice. **f,** Curve representing glycaemia (glucose levels) during glucose tolerance test in females 5 weeks after *AAV-Dio2-GFP* or *AAV-Dio2-Cre-GFP* injection and area under the curve (AUC). **g,** Representative infrared thermal images; **h,** eye temperature and **i,** BAT temperature 6 weeks after induction of the tanycytic cilia impairment in female mice. **j,** Relative expression of *Pgc1α*, *Tfam*, *Gk*, *Fasn*, and *Ucp3* in the BAT. **k,** Western blot and quantification of UCP1 protein levels, relative to the *Ift88^fl/fl^* group, in the BAT 6 weeks after induction of the tanycytic cilia impairment in female mice. All data are expressed as mean ± s.e.m., ∗, *p* < 0.05; ∗∗, *p* < 0.01; ∗∗∗, *p* < 0.001. *n* denotes the number of mice. Detailed information on the test statistics is provided in **Extended Data Table 4**.

Female mice with impaired tanycytic cilia displayed increased body weight after seven weeks (**Fig. 4d,e**) and showed a tendency toward glucose intolerance (**Fig. 4f**), whereas food intake and energy expenditure remained unchanged (**Extended Data Fig. 4a-c**). Tanycytic cilia impairment reduced core body temperature (**Fig. 4g,h**), as reflected by the non-invasive measurement of eye temperature^59,60^. Although the BAT showed only a non-significant trend toward lower surface temperature (**Fig. 4g,i**), the expression of key genes involved in BAT activity decreased at the transcript and protein levels, as shown by lower mRNA levels of *Tfam*, *Gk* and *Fasn* (**Fig. 4j**), as well as reduced UCP1 protein levels (**Fig. 4k**), following impairment of ciliary function.

In male mice, we observed the same IFT88 downregulation in tanycytes as in females (**Extended Data Fig. 3a-c**). However, the male mice with impaired tanycytic cilia did not show any differences in body weight, food intake, energy expenditure, core body and BAT temperatures (**Extended Data Fig. 4d-k**). This normal metabolic phenotype in male mice suggests a sexual dimorphism in tanycytic cilia functions. Together, the data indicate that even a partial loss of the ciliary protein IFT88 in tanycytes is sufficient to induce metabolic alterations in a sexually dimorphic manner, supporting a role for tanycytic cilia in maintaining metabolic homeostasis.

## Discussion

In this study, we identified tanycytic cilia as dynamic, metabolically sensitive organelles that integrate extracellular nutritional and hormonal cues with intracellular signalling responses in tanycytes. We show that tanycytic cilia exhibit subtype-, sex- and age-specific morphological features, are modulated by nutrient availability and thyroid hormones, and are required for proper Ca²⁺ signalling. Disrupting tanycytic cilia impairs ATP- and glucose-evoked Ca²⁺ responses, alters cAMP dynamics, impairs the autophagic flux and results in lipid droplet accumulation in tanycytes. Importantly, even a partial reduction in the intraflagellar transport protein IFT88 within tanycytic cilia in vivo was sufficient to promote metabolic dysregulation in female mice, including increased body weight, reduced core temperature, and decreased BAT activity (**Extended Data Fig. 5**). Together, these findings reveal tanycytic cilia as central regulators of tanycyte metabolic function and highlight their contribution to whole-body energy homeostasis.

Our analysis of tanycytic cilia reveals several layers of structural and functional specialization across tanycyte subtypes. We observed mainly uni-ciliated tanycytes. In the dorsal region, several α-tanycytes possessed two basal bodies as reported by Mirzadeh et al.^27^, but we found little evidence for the presence of two cilia on one cell when staining the tight junction protein ZO-1 to delineate tanycytic cell borders.

Subtype-specific differences in ciliary morphology suggest that tanycytes may use distinct ciliary structures to carry out their specialized functions. Consistent with such specialization, we found that in mice AC3, a marker of neuronal cilia^30^, is enriched in the β-tanycytes which are most capable of neurogenesis^61–63^, supporting the idea that these tanycytes share “neuronal-like” properties, at least in mice, and rely on cAMP-dependent signalling pathways associated with progenitor function.

We detected sex differences in cilia gene expression and in cilia length, especially in the ME, suggesting that tanycytic cilia may be modulated by sex hormones. Further supporting this theory, we found that tanycytic cilia morphology changes during the oestrous cycle. Whether these differences reflect direct hormone receptor signalling, circulating steroid fluctuations, or region-specific structural plasticity remains to be determined, but our observations raise the intriguing possibility that tanycytes integrate sex-specific metabolic or endocrine cues through ciliary architecture.

Aging alters primary cilia throughout the brain^11,31^, and our results show that tanycytes follow this pattern of age-related ciliary remodelling. We observed an elongation of tanycytic cilia with age, together with a selective loss of cilia in α1-tanycytes, which bear the longest cilia. Comparable elongation has been reported in hippocampal neurons^11,31^, where longer cilia correlated with reduced osteocalcin signalling and impaired cilia-dependent pathways. In contrast, *Mc4r*-expressing hypothalamic neurons exhibit age-related cilia shortening that diminishes satiety signalling and contributes to metabolic dysfunction^13^. These findings illustrate that aging does not uniformly impact all cilia but rather perturbs the cell-type-specific morphology required for optimal sensory function. Because both overly long and shortened cilia disrupt signaling^64–66^, the age-related elongation and in particular the loss of cilia that we observed in old mice may ultimately compromise tanycyte function. Together, these observations support the emerging concept of an age-related ciliopathy^13^, in which altered cilia structure contributes to impaired metabolic communication and increases vulnerability to energy imbalance during aging.

TSH treatment induced a notable increase in ciliary length, consistent with recent findings from Melum and colleagues^26^, suggesting that TSH from the pars tuberalis (PT) modulates the expression of ciliary genes in tanycytes. In their photoperiodic model, long photoperiods, associated with elevated PT-TSH, upregulated ciliary assembly-related genes. The metabolic effects of tanycytic cilia that we have discovered would suggest that TSH adapts metabolism to seasonal demands by controlling the function of tanycytic cilia.

In contrast to serum-starvation, which promoted ciliogenesis in primary tanycytes, oleic acid induced ciliary shortening. Links between fatty acid exposure, ciliary signalling, and metabolic reprogramming have been established in hypothalamic neurons, where impaired ciliary function disrupts autophagy and promotes metabolic remodeling^39,67^. The primary cilium in kidney epithelial cells is responsible for mediating similar changes in autophagy^53^. Since lipophagy and ciliary function can be mutually dependent^68^, the question arises whether ciliary changes in oleic acid-treated cells are the cause or consequence of less lipophagy. Our observation that *Ift88* knockdown led to lipid droplet accumulation suggests that oleic acid-induced tanycytic cilia shortening impairs lipophagy, resulting in lipid droplet accumulation and metabolic reprogramming under energy-surplus conditions. In tanycytes, lipophagy regulates lipid droplet homeostasis and fuels ATP production^45^.

Given that tanycytes detect blood glucose levels at the ME and convert them into propagating Ca²⁺ waves, we examined whether tanycytic cilia contribute to glucose- and ATP-evoked Ca²⁺ signalling. In primary tanycytes with impaired cilia, Ca²⁺ responses to both glucose and ATP were diminished, indicating that functional cilia amplify these signals. Together, these results reveal that tanycytic cilia are not passive structural appendages but rather central hubs orchestrating Ca²⁺ and cAMP signalling in tanycytes. When ciliary integrity is compromised, either genetically or through metabolic overload, tanycytes lose their ability to decode ATP and glucose cues, accumulate lipids, and exhibit impaired autophagy. This positions tanycytic cilia as essential organelles that couple extracellular metabolic information to intracellular metabolic state and suggests that ciliary dysfunction may represent a critical vulnerability linking nutrient excess to tanycyte dysregulation. However, because our in vitro stimulation reached both apical cilia and basal processes, the polarity of glucose sensing, blood-derived glucose via tanycyte processes in the ME versus CSF-derived glucose or ATP at the apical surface, could not be disentangled. The residual Ca²⁺ response following ciliary knockdown suggests that non-ciliary glucose-sensing pathways remain active. Nonetheless, our findings support a model in which tanycytic cilia contribute to glucose sensing, possibly by detecting glucose-derived ATP in the CSF.

We found that impairment of tanycytic cilia, in vivo, led to an increase in body weight, a tendency toward glucose intolerance, reduced core body temperature, and decreased BAT activity, highlighting a broad impact on systemic metabolic regulation. We did not observe changes in cilia number in vivo following *Ift88* knockdown, which may reflect the timing of gene disruption in adult mice, when cilia number is largely stabilized. This underscores the previously known dispensability of *Ift88* in cilia structure retention and highlights its role for the maintenance of sensory cilia function^51^. Compensatory mechanisms might also contribute to preserving ciliary abundance despite partial impairment. Interestingly, even a partial reduction in *Ift88* expression was sufficient to alter metabolic parameters, strongly implicating tanycytic cilia in metabolic regulation. The metabolic phenotype that we observed after tanycytic cilia impairment was specific to female mice, in line with the sexual dimorphism in the length of tanycytic cilia and ciliary gene expression. One possibility is that the shorter length of tanycytic cilia makes females more vulnerable to the metabolic consequences of ciliary dysfunction and ultimately to obesity^69^.

Together, our findings position tanycytic cilia as dynamic sensory organelles capable of integrating hormonal, metabolic, and environmental cues. Their structural plasticity and subtype-specific specialization underscore their potential central role in coordinating hypothalamic function across different physiological states.

## Material and methods

### Human tissue

Human hypothalamic tissue samples were processed for immunofluorescence staining following standardized neuropathological protocols. Tissue was fixed in 4 % phosphate-buffered formaldehyde, embedded in paraffin, and sectioned at 30 µm thickness using a microtome. Tanycyte and cilia markers (vimentin, ARL13B) were visualized using host-specific primary antibodies and species-appropriate Alexa Fluor secondary antibodies (AF488, AF568). TrueBlack was applied to quench lipofuscin autofluorescence. Images were acquired using an Olympus FV1000 laser scanning microscope with a 63x objective.

### Single-cell/nuclei sequencing data

Single-cell RNA sequencing from the mouse and single-nuclei RNA-sequencing from the human mediobasal hypothalamus were used to analyse the expression of ciliary genes in tanycytes^28^. Briefly, the four tanycytic subtypes were subset based on known markers and were reclustered (dimensions 1:28, resolution = 0.6) into 8 subclusters. Single-cell data for intraflagellar transport and cilium assembly markers were retrieved from Tanybase (https://data.mendeley.com/datasets/p6jkzkpdd6/2; github https://github.com/umitozorhan/Tanybase). Gene expression was visualized using DotPlot() in Seurat R package (v.5.3.0).

### Animals

#### Mouse lines

The following strains were used: *Ift88^fl/fl^* (RRID:IMSR_JAX:022409) previously described in^57,58^. All lines were maintained on a C57BL/6 background. Wild-type C57BL/6 mice were obtained from Charles River Laboratories (Sulzberg, Germany). Animals were housed at 22 °C on a 12-h light/dark cycle with ad libitum access to food (Altromin #1314) and water.

#### Study approvals

All animal experiments were conducted in accordance with EU Directive 2010/63/EU and approved by the local regulatory authority (Ministerium für Landwirtschaft, ländliche Räume, Europa und Verbraucherschutz, approval number 9-2/23). Human brain samples used in this study were obtained from body donations received by the Institute of Anatomy in Leipzig. The use of human brain tissue from body donations was approved by the Ethics Committee of the University of Leipzig (129-21 ek). All donors or their relatives provided written informed consent. All tissue retrievals were in line with the Saxonian Death and Funeral Act of 1994, third section, paragraph 18, item 8.

##### AAV vector production and stereotactic injection

To transduce tanycytes in vivo, we produced the AAV vectors *AAV-Dio2-Cre-2A-GFP* and *AAV-Dio2-GFP* with a mosaic capsid of serotypes 1 and 2^24^. After purifying vectors by an iodixanol gradient as described previously^70^, we measured the viral titre by real-time PCR with primers directed against the WPRE (**Extended Data Table 1**). For the stereotactic injection, mice were anesthetized with ketamine (30 µg/g body weight, i.p.) and isoflurane (induction, 3%; maintenance, 1%). After fixing mice in a stereotactic frame (Kopf Instruments), AAVs (5 x 10^9^ gp in 2 µl) were injected into the lateral ventricle as described previously^24^.

### Physiological measurements

#### Indirect calorimetry

Total energy expenditure, oxygen consumption (VO₂), CO2 production (VCO₂), respiratory exchange ratio (RER), ambulatory activity, and food intake were monitored using TSE Systems calorimetry cages. Mice were acclimated to cages for 48 h prior to data collection. RER and energy expenditure (EE) were calculated as reported previously^71,72^.

#### Glucose tolerance test

Following a 6-hour fasting period, basal blood glucose levels were measured before and 15, 30, 60 and 120 min after an intraperitoneal injection of D-glucose (Sigma-Aldrich, G8270) 2 g/kg body weight, with a glucometer (Accu-Chek Performa, Roche). Area under the curve (AUC) values were determined.

#### Thermography

The eyes and interscapular temperatures were measured three times using a high-resolution infrared camera (VarioCAM® hr head; InfraTec) and analysed with the IRBIS 3.0 software package.

### Brain and peripheral tissue processing

#### Immunofluorescence staining

Mice were deeply anesthetized and perfused with 4% PFA. Tissues were postfixed with 4% PFA for 16 h at 4 °C. Then, tissues were embedded in 4% low melting agarose and coronal 50-μm thick brain sections were cut with a vibratome (Leica VT1200S). If not specified otherwise, we investigated sections containing the ME (zones 2 and 3, **Extended Data Fig. 1d**). Antigen retrieval with sodium citrate buffer (10 mM sodium citrate, 0,05% Tween20, pH = 6) at 95 °C for 10 min was performed, followed by 3% H_2_O_2_ in methanol for 15 min prior to blocking, permeabilized and blocked for 2 h in 0.3% Triton X-100 and 5% normal goat or donkey serum in PBS at room temperature and then incubated overnight at 4 °C with primary antibodies (**Extended Data Table 2**) diluted in blocking solution. Sections were washed with PBS containing 0.3% Triton X-100 before and after being incubated for 2 h with secondary antibodies (**Extended Data Table 2**) at room temperature in blocking solution containing DAPI to stain nuclei (1 µg/ml, Thermo Fisher). All sections were mounted onto SuperFrost Plus slides using fluorescence mounting medium (DAKO) and let dry over night before imaging.

#### Imaging

Confocal microscopy was performed using a Leica TCS SP5 or Stellaris 5 microscope equipped with a 63X immersion (oil) objective. Images were acquired using LAS X (Leica) software and processed with Imaris (Oxford Instruments). A custom-built STED microscope (Abberior GmbH) was used for super-resolution microscopy. The setup included a 560- or 640-nm diode excitation laser, a 775-nm STED laser, both pulsed at 40 MHz, and a 1.4 NA objective (UPlanSApo 100x/1.40 oil, Olympus). A z-scanning piezo (PIFOC, Physik Instrumente) was used for focusing, and a spatial light modulator (Hamamatsu) was used in the depletion beam to generate a top-hat (3D-STED) phase mask. Sample induced aberrations were corrected using a deformable mirror. For Dynamic Intensity Minimum (DyMIN) STED, the 3D-STED imaging was performed with 80% 3D-STED beam. Images were recorded using 40 × 40 × 40 nm voxel size, and the DyMIN threshold was set to 5 counts. The exclusion parameter was set to 5. Images acquired with the STED microscope were first deconvolved using the Richardson-Lucy algorithm with the “DeconvolutionLab2” plugin (Fiji) and then drift-corrected with the StackReg plugin (Fiji).

#### Cilia number and length

To assess cilia number and length, cilia were traced in 3D using the Imaris Filament Tracer module (Oxford Instruments, Abingdon, UK). The basal body, stained with pericentrin, served as a reference to identify the origin of the ciliary axoneme. The axoneme, labelled with ARL13B, was traced from base to tip using the Pencil tool in Auto Path mode. Imaris automatically centred each traced cilium once it was manually delineated. Cilia number was normalised to the volume of the tanycytic DAPI surface, generated with the Imaris Surface module. Surfaces defining tanycytic subpopulations were manually drawn based on the classification of tanycytes into the four zones of the 3V^73^. Imaris statistics, including cilia length, number of cilia per image, and tanycytic DAPI volume, were exported to Excel for further analysis. Final measurements included ciliary length (µm) and the ratio of cilia per volume for each tanycytic subpopulation, referred to in the graphs as *“number of cilia normalized to ROI.”*

#### AC3/ARL13B colocalization analysis

To assess colocalization between AC3 and ARL13B, mask selections of the previously generated surfaces delimiting the respective tanycytic subpopulation were created. ARL13B and AC3 signals were segmented in their respective channels. Colocalization was quantified using the *“Shortest Distance to ARL13B Surface”* filter, with a threshold set to ≤ 0. The average volume of AC3 surfaces within a distance ≤ 0 to the ARL13B surface, as well as the average volume of the ARL13B surface, were exported to Excel files. The ratio of AC3-positive to total ARL13B surface volume was then calculated.

#### Ciliary IFT88 analysis

To assess the number of IFT88 puncta in ARL13B positive cilia, mask selections of the previously generated surfaces delimiting the respective tanycytic subpopulation were created. To assess IFT88 and ARL13B signals, the spot detector and surface tools were used respectively. Selection of the ciliary IFT88 puncta was quantified using the *“Shortest Distance to ARL13B Surface”* filter, with a threshold set to ≤ 0. The number of IFT88 spots within a distance ≤ 0 to the ARL13B surface, as well as the average volume of the ARL13B surface, were exported to Excel files. The ratio of the number of ciliary IFT88 spots to the nuclei volume is normalized to the control group.

#### RNA extraction from organs and quantitative RT-PCR

BAT tissues were snap-frozen and homogenized with an Ultra Turrax homogenizer. RNA was extracted using the NucleoSpin RNA kit (Macherey-Nagel). Then, mRNA (400 ng) was reverse transcribed using the M-MLV reverse transcriptase and random primers. Primers for real-time PCR are listed in **Extended Data Table 1**. Platinum SYBR Green qPCR SuperMix (Invitrogen) and the ROCHE Light cycler were used with the following protocol: 10 min at 95 °C preincubation), 30 s at 95 °C, 30 s at 60 °C and 30 s at 72 °C (3-step amplification with 38 cycles). *Ppia* served as housekeeping gene for data normalization according to the ΔΔCt method.

#### Immunoblot analyses

Tissues were homogenized using a TissueLyser II (Qiagen) in cold RIPA buffer (200 mM Tris/HCl pH 7.4, 130 mM NaCl, 10% (v/v) glycerol, 0.1% (v/v) SDS, 1% (v/v) Triton X-100, 10 mM MgCl_2_) with anti-proteases and anti-phosphatases (Sigma-Aldrich), and lysates were centrifuged for 30 min at 18,000 x *g* and 4 °C. BAT total protein lysates were separated on SDS–polyacrylamide gel electrophoresis (SDS–PAGE), electrotransferred to nitrocellulose membranes and probed successively with primary antibodies (listed in **Extended Data Table 2**) after incubation of membranes with 5% BSA blocking buffer. Proteins were detected using horseradish peroxidase–conjugated secondary antibodies (**Extended Data Table 2**). Specific immunolabelling was visualized using chemiluminescence following the manufactureŕs instructions (Pierce ECL Western Blotting Substrate, ThermoScientific), and values expressed relative Actin or Tubulin. Primary tanycytes experiments

#### Culture of primary tanycytes

Primary tanycytes were extracted from P10 rats by carefully dissecting the wall of the 3V and the ME as previously outlined^74,75^. In short, after rinsing the brains three times with ice-cold PBS, the MBH and ME were separated under a binocular microscope. The tissue was placed in an ice-cold growth medium (DMEM with high glucose (Thermo Fisher Scientific #41965), 10% donor bovine serum (DBS), 1% penicillin/streptomycin, and 2 mM l-glutamine). The tissue fragments were then passed through a 70-µm nylon mesh (Merck Millipore) and centrifuged at 440 x *g* (8 min at 4 °C). After resuspending the cells in fresh growth medium, they were seeded into T75 cell culture flasks (Cellstar, Greiner Bio-one). Ten days after preparation, the medium was changed twice weekly, until the cells achieved confluence. The cultures typically consisted of over 80% vimentin-positive tanycytes. The primary tanycytes were then split using accutase (Sigma-Aldrich) and replated into 12-, or 24-well plates at a density of 2 × 10⁵ or 1 × 10⁵ cells per well, depending on experimental requirements.

#### Starvation

Primary tanycytes were incubated for 16 h in prewarmed starvation medium (DMEM/F-12, HEPES, without phenol red; Thermo Fisher Scientific #11039), supplemented with 0.02% putrescine, 0.01% insulin, and 1% penicillin/streptomycin. Control cells were maintained under identical conditions in prewarmed full culture medium.

#### Free fatty acids treatments

Primary tanycytes were treated with 100 µM OA (Sigma-Aldrich, #O1008), or 100 µM PA (Sigma-Aldrich, #P0500) conjugated to fatty acid-free bovine serum albumin (BSA), or a mix of both for 6 h. Vehicle-treated cells received the same percentage of methanol (0.25%) and/or BSA (0.06%), diluted in culture medium. The concentration of PA and OA were chosen as previously described to mimic the levels of fatty acids in the brain of a long-term diet-induced obese mouse^39,76^.

#### Thyroid hormone treatments

Primary tanycytes were starved in starvation medium for 6 h and then incubated with either 10 nM T3 (Sigma-Aldrich, T23877), 10 nM T4 (Sigma-Aldrich, T2376), or 60 mIU TSH (Sigma-Aldrich, T8931)) for 16 h in thyroid hormone depleted culture medium, as previously described^34^. Control cells were also starved for 6 h and then received 500 µL of prewarmed (37 °C) thyroid hormone–depleted culture medium (DMEM with high glucose (Thermo Fisher Scientific #41965) supplemented with 10% charcoal stripped foetal bovine serum (Thermo Fisher Scientific, 12676029), 1% penicillin/streptomycin (100×; Biochrom, A2213), and 2mM L-glutamine) per 24-well.

#### Autophagy inhibition

Primary tanycytes were treated with Bafilomycin A1 (BafA1; B1793, Sigma-Aldrich) at a concentration of 100 nM for 4 h before fixation or protein lysis. DMSO (BM-0660, Winkler) treated cells were used as controls for BafA1.

#### siRNA Transfections

Cells were transfected two days after splitting, after 4 hrs incubation in antibiotic-free medium, using Lipofectamine™ RNAiMAX (Thermo Fisher Scientific, Invitrogen, Waltham, MA, USA) according to the manufacturer’s instructions. Small interfering ribonucleic acids (siRNAs) targeting *Kif3A* and *Ift88* mRNAs were used, while a non-targeting siRNA pool (scramble) served as control (Dharmacon: ON-TARGETplus siRNA-*Ift88* (rat): CGUUGGAAAUUGACGAAGA; AGCAAAUGUAUACGAGUUA; CUGGGAGAGUUGUACGAUA; GCUAUUAAAUUCUACCGAA. ON-TARGETplus siRNA-*Kif3a* (rat): GAUGGAAAUGCAAGCGAAA; CUGACGACAUGGAUAGAAU; AAUCGAGUGCAGCGAGAAA; GCAAAGCCUGAGACCGUAA. ON-TARGETplus Non-targeting Pool (Scramble): Negative control siRNA with at least 4 mismatches to rat gene). After transfection, cells were returned to the incubator and maintained for 24–72 h before further treatments.

#### RNA isolation from primary tanycytes

RNA was isolated from primary tanycytes using the NucleoSpin RNA kit (Macherey-Nagel). Then, mRNA (150 ng) was reverse transcribed using the M-MLV reverse transcriptase and random primers. Primers for real-time PCR are listed in **Extended Data Table 1**. Platinum SYBR Green qPCR SuperMix (Invitrogen) and the ROCHE Light cycler were used with the following protocol: 10 min at 95 °C preincubation), 30 s at 95 °C, 30 s at 60 °C and 30 s at 72 °C (3-step amplification with 38 cycles). *Ppia* served as housekeeping gene for data normalization according to the ΔΔCt method.

#### Western blotting

For western blotting, cells were washed with PBS and then collected in 150 µl lysis buffer while shaking (50 mM Tris-HCl pH 7.5, 1 mM EGTA, 1 mM EDTA, 1% Triton X-100, 1 mM sodium orthovanadate, 50 mM sodium fluoride, 5 mM sodium pyrophosphate, 0.27 M sucrose, 1 mM phenylmethylsulfonyl fluoride (Carl Roth 6367.2) and 1x protease inhibitor (cOmplete, Roche 11836153001). Cell lysates were centrifuged at 18,620 x *g* for 5 min at 4 °C. The supernatant was mixed with SDS buffer (0.75 M Tris-HCl, 0.08 g/ml SDS, 40% glycerol, 0.4 mg/ml bromophenol blue, and 62 mg/ml DTT) in a 3:1 ratio. After samples were subjected to SDS-PAGE, the proteins were transferred to nitrocellulose membranes. Membranes were incubated with primary antibodies (**Extended Data Table 2**) diluted in blocking solution overnight at 4 °C and the respective HRP-conjugated secondary antibodies (**Extended Data Table 2**) for 2 h at RT. Finally, we detected the signal using enhanced chemiluminescence (SuperSignal West Pico Substrate, Thermo Scientific, 34580) and a digital detection system (Fusion Solo S, Viber). Protein quantification was performed using ImageJ software, and protein levels were normalised to Actin. Autophagic flux was calculated as the difference in LC3II levels in presence and absence of BafA1 (100 nM).

#### Immunohistochemistry

Cells were seeded on poly-D-lysine-coated cover slips (1 x 10^5^ per well in a 24-well plate). Following treatment, cells were fixed with 4% PFA for 15 min at RT, washed with PBS 1x. Fixed cells were permeabilized with 0.3% Triton X-100 in PBS (1x) for 15 min, followed by blocking in 5% bovine serum albumin (BSA) in 0.3% Triton X-100/PBS for 1 h at RT. Cells were then incubated overnight at 4 °C with primary antibodies diluted in the same blocking solution (**Extended Data Table 2**). The next day, cells were incubated for 1 h at room temperature with appropriate secondary antibodies, also diluted in blocking solution (**Extended Data Table 2**). For lipid droplets and autophagic puncta visualization, no triton was used but 0,5% Saponin was used as unique permeabilization. Nuclei were labelled with DAPI. Coverslips were mounted onto SuperFrost Plus slides using fluorescence mounting medium (DAKO) or an aqueous mounting medium for lipid droplets dye and let dry over night before imaging.

#### Imaging

Confocal microscopy was performed using a Leica TCS SP5 microscope equipped with a 63X immersion (oil) objective. Images were acquired using LAS X (Leica) software.

#### Identification of primary tanycytes

Primary tanycytes were identified based on nuclear morphology in the DAPI channel and cell body morphology in the vimentin channel. Only cells meeting the predefined criteria for primary tanycytes were included in the analysis. These criteria comprised a large, oval-shaped nucleus and an intense, spread vimentin staining pattern.

#### Cilia analysis

Cilia length, cilia number, and nuclei per image were quantified using Imaris software (Oxford Instruments). Cilia were traced using the Filament Tracer module (diameter: 0.36 µm), and nuclei were identified using the Spot Detector module (diameter: 6 µm). The number of cilia was normalized to the number of DAPI-positive nuclei. Nuclei located at the image border were excluded from analysis if they could not be reliably assigned to a cilium.

#### Lipid droplet and autophagy puncta analysis

ImageJ software 1.54p was used to quantify the LC3, p62 and Lipidye puncta (Analyse particle, size (µm²): 0.01; 0.1; 0.1 respectively) as well as Dapi number to assess the cell number (Analyse particle, size (µm²): 50) and Vimentin area (Analyse particle, size (µm²): 0.0001). For colocalization analysis between LC3 and Lipidye puncta, the “Coloc2” function of ImageJ was used to generate Pearson’s correlation coefficient values which ranged from −1 (perfect exclusion) to +1 (perfect correlation).

### Functional assays

#### Intracellular calcium measurements

Cells were seeded on poly-D-lysine-coated cover slips (1 x 10^5^ per well in a 24-well plate) and incubated in Fluo4-AM solution (2 µM Fluo4-AM, 0.2 mg/mL pluronic 127, 2.5 mM probenecid in 1.5 ml medium) for 30 min at 37 °C and 5% CO2. The coverslips were then placed in a flow chamber that was perfused with artificial cerebrospinal fluid (aCSF; 12.4 mM NaCl, 2.6 mM NaHCO_3_, 0.125 mM NaH_2_PO_4_, 0.3 mM KCl, 0.1 mM MgSO_4_, 0.2 mM CaCl_2_, 5% CO2/ 95% O2) containing 1 mM glucose and 9 mM sucrose. After 1 min of baseline recording, primary tanycytes were stimulated with one of the following compounds: 20 mM D-glucose (Sigma-Aldrich), 33 µM taltirelin (Tocris Bioscience) or 333 µM ATP (Sigma-Aldrich). For the specific case of glucose stimulation, cells were previously incubated for 48 h in a low-glucose medium (D-glucose 2mM) before calcium imaging. Fluorescence was imaged for 5 min at 1 image/sec (300 images), using a setup (Till Photonics) mounted on the Axio Examiner D1 upright fluorescence microscope (Zeiss). For analysis, labelled tanycytes were marked as regions of interest (ROI) and the fluorescence intensity of each ROI was normalized to baseline (30 sec before stimulation). Cells were only included in the analysis, if the fluorescence intensity peaked within 60 sec after stimulation and then returned to baseline levels. For each coverslip, 50 ROIs were manually drawn and analysed using ImageJ software (NIH, Bethesda, MD, USA).

#### Intracellular cAMP measurements

For the cAMP-Glo assay (Promega), primary tanycytes were seeded on 96-well plates (0.4 x 10^5^ cells per well). When the cells reached approximately 80% confluence and after 72 h of transfection, the medium was removed and the tanycytes were stimulated for 30 min with forskolin (25 µM, Sigma-Aldrich, F3917) dissolved in induction buffer. Controls were treated with the induction buffer only. Cells were then washed with PBS and lysed in lysis buffer. cAMP-Glo detection solution (40 µl provided by the kit) was added to the cell lysate and incubated for 20 min at room temperature (RT). After the addition of Kinase-Glo (80 µl) and incubation for 10 min at RT, luminescence was measured using the Clario Star Plate Reader (BMG Labtech) with a detection wavelength range between 320 and 700 nm. Raw luminescence values were converted to cAMP levels using the formula cAMP = 1/luminescence_raw_. For final analysis, cAMP levels after forskolin stimulation were normalised to their respective controls (e.g., Scramble + forskolin was normalised to Scramble basal).

#### Statistical data analysis

Statistical analyses and graphical representations were prepared using GraphPad Prism 8 (GraphPad Software) and SPSS Version 29 (IBM). All values are presented as mean ± SEM (standard error of the mean). Statistical tests and parameters, including sample size (n), post hoc tests, and exact p-values, are reported in **Extended Data Table 4**. Parametric statistics (e.g., t-Test, ANOVA) were only applied if assumptions were met, i.e., datasets were examined for Gaussian distribution by D’Agostino-Pearson test, aided by visual inspection of the data, and homogeneity of variances by Brown-Forsythe, Levene’s or F-test (depending on the statistical method used). If assumptions for parametric procedures were not met or could not be reliably assumed due to small sample size, non-parametric methods were used as indicated. Two-tailed tests were applied if not indicated otherwise. Greenhouse-Geisser correction was used in ANOVA statistics if the sphericity assumption was violated (Mauchly test). Sample sizes were estimated based on pilot experiments. When applicable, an outlier detection method (ROUT method, Q = 1%) was used. In figures, statistical significance is indicated as follows: *, *p* < 0.05; **, *p* < 0.01; ***, *p* < 0.001.

## Acknowledgements

Special thanks go to Ines Stölting and Frauke Spiecker for excellent experimental support. This work was supported by grants from the European Research Council (Synergy grant no. 2019-WATCH-810331 to R.N., V.P. and M.S.; HORIZON TMA MSCA Postdoctoral Fellowships no. 101154613 – Tanycilia to M.R.).

## Author contributions

M.R. and M.S. conceived the project. M.R., C.K. and M.S. designed experiments. M.R., R.N., V.P. and M.S. procured funding. M.R., C.K., A.C., W.B., S.D., S.G., Ü.Ö., S.N., J.R., N.F., V.N., S.R., S.B. and H.M.F. performed experiments and analysed data. I.B., R.N., V.P. and M.S. provided unique reagents, infrastructure, mouse models and viral vectors. M.R., C.K. and M.S. wrote the manuscript with input from all co-authors.

## Competing interests

The authors declare no competing interests.

## Notes

### Competing Interest Statement

The authors have declared no competing interest.

