## Supplementary figures and tables for "Ciliary sensing in tanycytes couples nutrient availability to metabolic regulation"

**The PDF file includes:**

Extended data figures 1 to 5

Extended data tables 1 to 4

### Extended data figures

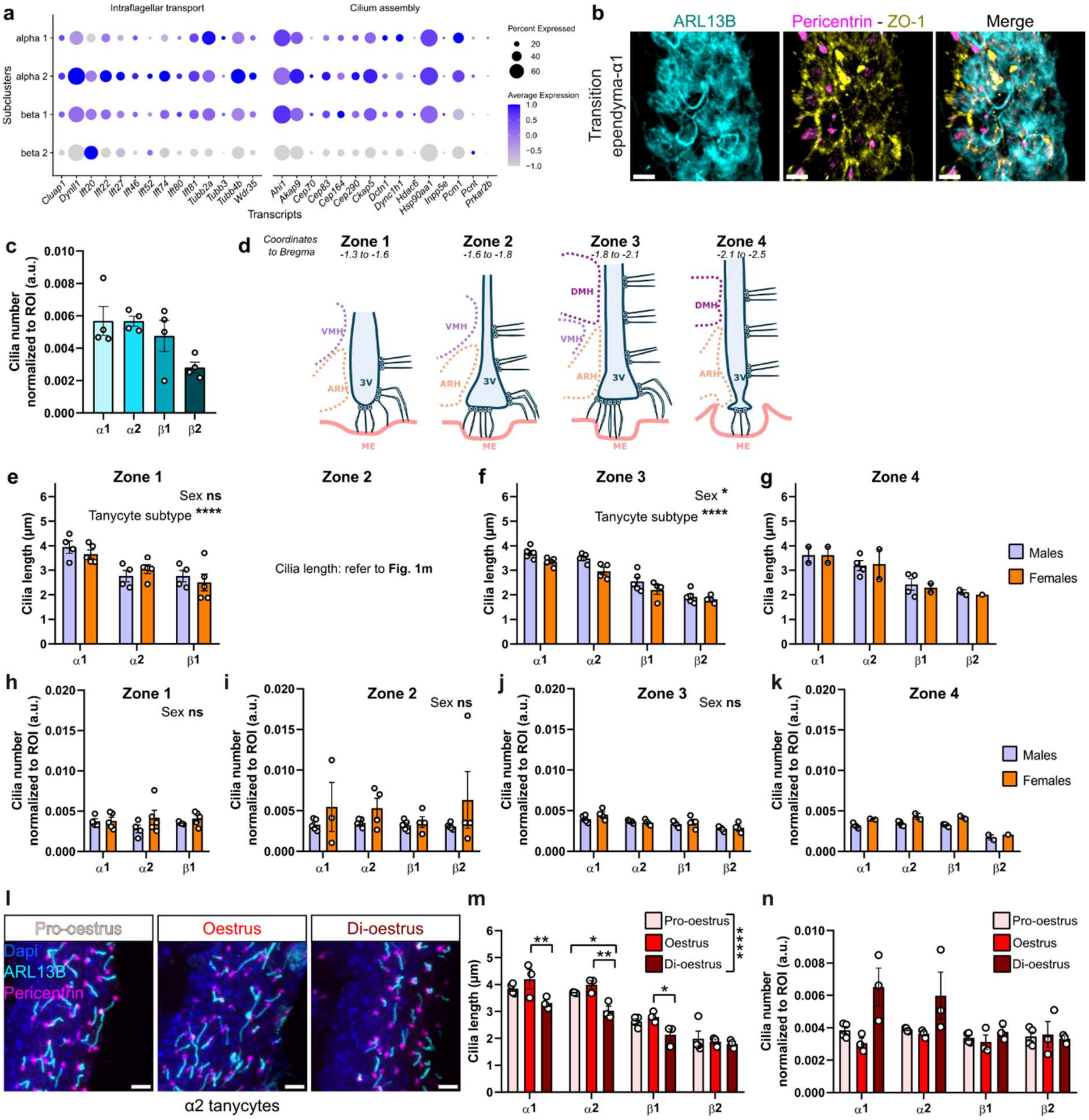

**Extended Data Fig. 1. Tanycilia display a subtype variability on the dorsoventral and anteroposterior axes and are modulated by sex hormones.**

**a,** Single-cell RNA sequencing data from Tanybase for intraflagellar transport and cilium assembly genes for the tanycyte subgroups, displaying the variability in ciliary gene expression among tanycyte subtypes. **b,** Immunofluorescent staining of cilia (ARL13B, cyan), basal body (pericentrin, magenta) and tight junctions (ZO-1, yellow) in a coronal section of a mouse brain, representative images of the ependyma-α1-tanycyte transition. Scale bar, 5 µm. **c,** Cilia number normalized to ROI in the 4 tanycyte subtypes in control mice. **d,** Scheme representing the 3V zones divided according to the antero-posterior axis. **e-k,** Cilia length and quantification of cilia number normalized to ROI in the 4 tanycyte subtypes from male and female control mice, in the tanycyte zones 1 **(e, h)**, 2 **(i)**, 3 **(f, j)** and 4 **(g, k)** according to the anteroposterior axis (Zone 2 and 3 are in the ME). **l,** Cilia and basal body immunofluorescent staining (ARL13B, cyan; pericentrin, magenta) in α2-tanycytes of pro-oestrous, oestrous and di-oestrous female mice. Scale bar, 5 μm. **m,n,** Cilia length **(m)** and quantification of cilia number normalized to ROI **(n)** in the 4 tanycyte subtypes from 4 pro-oestrous, 3 oestrous and 3 di-oestrous female mice. All data are expressed as mean ± s.e.m., ∗, $p < 0.05$; ∗∗, $p < 0.01$; ∗∗∗, $p < 0.001$. *n* denotes the number of mice. Detailed information on the test statistics is provided in **Extended Data Table 4**.

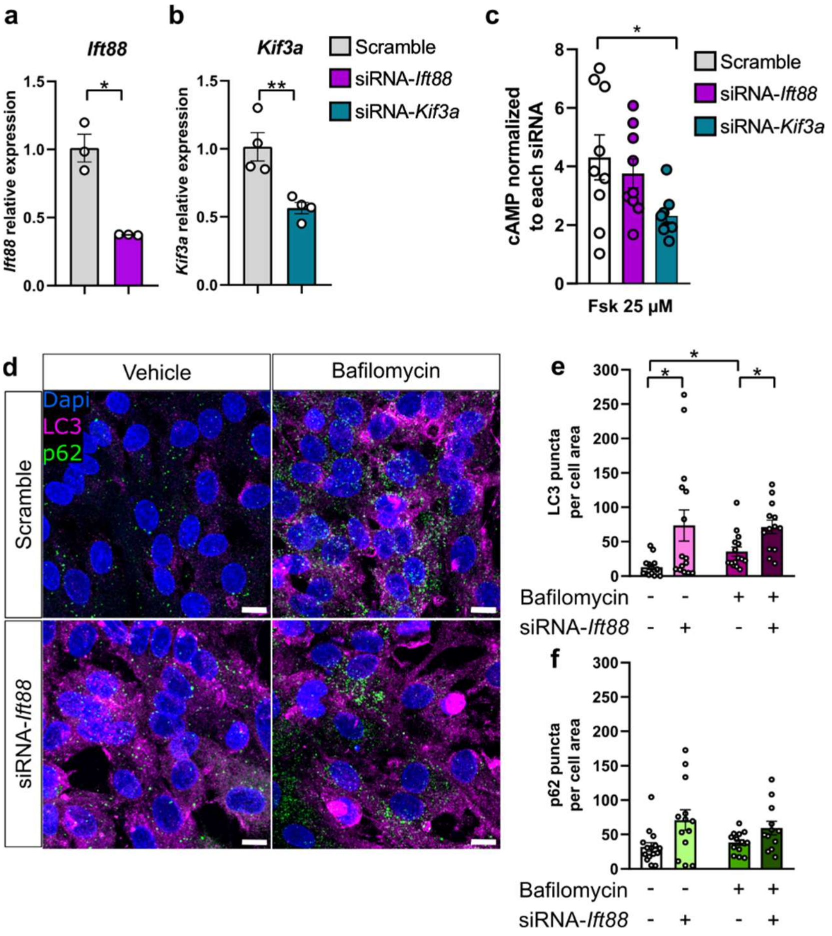

**Extended Data Fig. 2. Ciliary impairment leads to decreased cAMP signalling and impaired autophagic flux.**

**a,b,** Relative expression of *Ift88* **(a)** and *Kif3a* **(b)** mRNA levels in primary tanycytes 72 h after transfection with siRNA-scramble, siRNA-*Ift88* or siRNA-*Kif3a*. **c,** cAMP levels in transfected primary tanycytes after 10 minutes of stimulation with 25 µM of forskolin. The values for the forskolin treated groups were obtained by normalising each group to its respective basal cAMP levels. **d,** LC3 and SQSTM1/p62 puncta stainings (LC3, magenta; p62 green) in primary tanycytes 72 h after transfection with siRNA-scramble or siRNA-*Ift88*, treated with DMSO of Bafilomycin (BafA1, 100nM) for 4 h (Scale bar, 10 µm), and **e** quantification of LC3 puncta per cell area and **f** p62 puncta per cell. All data are expressed as mean ± s.e.m., ∗, $p < 0.05$; ∗∗, $p < 0.01$; ∗∗∗, $p < 0.001$. *n* denotes the number of wells **(a-c)** or single images analysed from 3 wells **(e,f)**. Detailed information on the test statistics is provided in **Extended Data Table 4**.

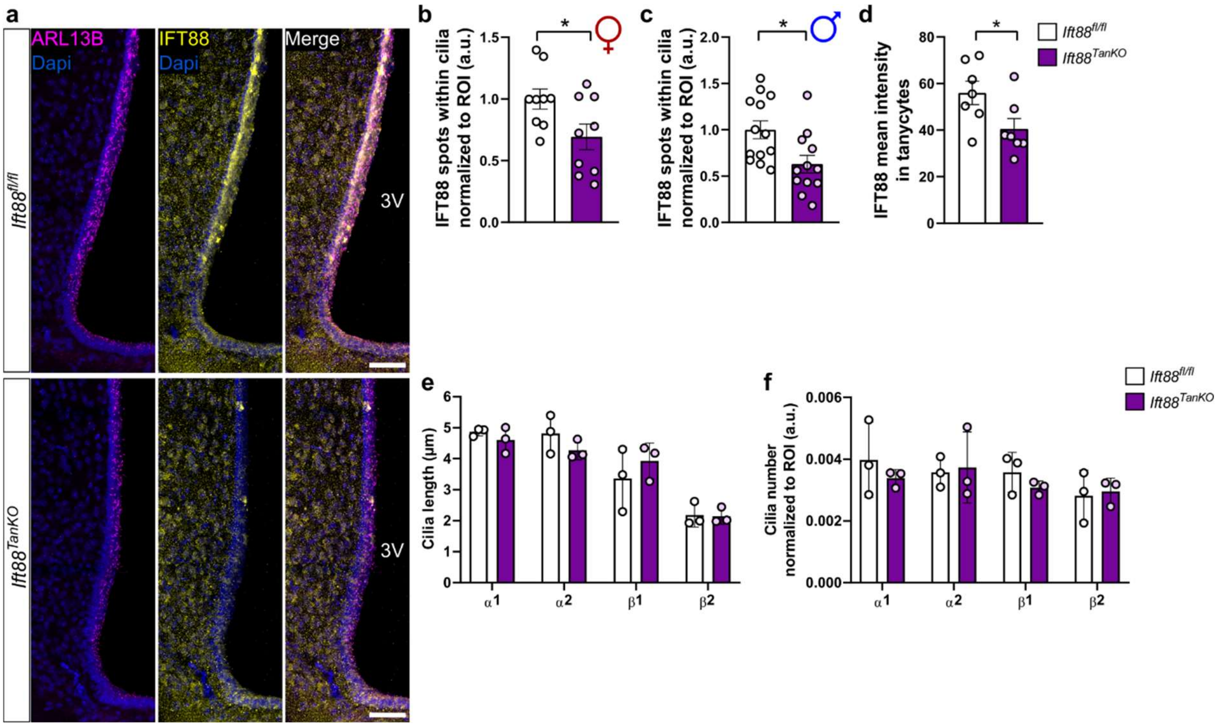

**Extended Data Fig. 3. Establishment of a tanycyte-specific impairment of cilia in an adult mouse model.**

**a,** Overview representative images of tanycytic cilia (ARL13B, magenta) and IFT88 (yellow) in coronal sections of the 3V, 4 weeks after injection of *AAV-Dio2-GFP* or *AAV-Dio2-Cre-GFP* in the lateral ventricle of *Ift88*$^{fl/fl}$ adult mice. Scale bar, 50 µm. **b,c,** Quantification of IFT88 spots within cilia per ROI (versus *AAV-Dio2-GFP*) in female **(b)** and male mice **(c)**. **d,** Quantification of IFT88 mean intensity in the tanycyte layer of both male and female mice. **e,f,** Cilia length measurement **(e)** and quantification of cilia number normalized to ROI **(f)** in the 4 tanycyte subtypes, 4 weeks after induction of the tanycytic cilia impairment in male mice. All data are expressed as mean ± s.e.m., ∗, $p < 0.05$; ∗∗, $p < 0.01$; ∗∗∗, $p < 0.001$. *n* denotes the number of mice. Detailed information on the test statistics is provided in **Extended Data Table 4**.

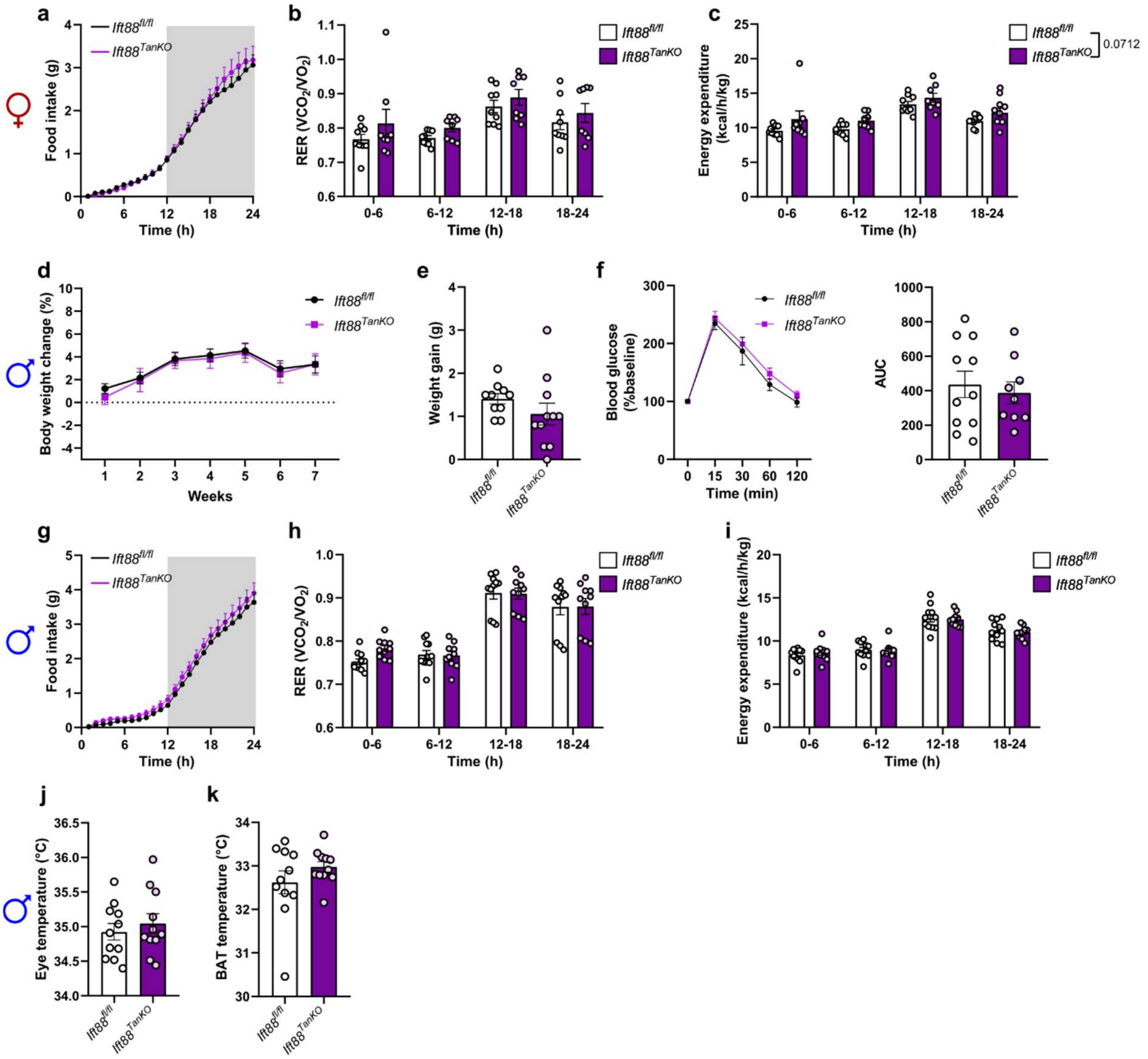

**Extended Data Fig. 4. Metabolic investigation in a tanycyte-specific impairment of cilia in an adult mouse model.**

**a,** Food intake over time, **b,** mean respiratory exchange ratio (RER) and **c,** mean energy expenditure during phases of 6 h in female mice, 7 weeks after induction of tanycytic cilia impairment. **d,** Body weight change (%) from the week of injection to 7 weeks after induction of the tanycytic cilia impairment in male mice. **e,** Weight gain 7 weeks after induction of the tanycytic cilia impairment in male mice. **f,** Curve representing glycaemia (glucose levels) during glucose tolerance test in male mice, 5 weeks after *AAV-Dio2-GFP* or *AAV-Dio2-Cre-GFP* injection and area under the curve (AUC). **g,** Food intake over time, **h,** mean respiratory exchange ratio (RER) and **i,** mean energy expenditure during phases of 6 h in male mice, 7 weeks after induction of tanycytic cilia impairment. **j,** eye temperature and **k,** BAT temperature obtained by infrared camera 6 weeks after induction of the tanycytic cilia impairment in male mice. All data are expressed as mean ± s.e.m., ∗, $p < 0.05$; ∗∗, $p < 0.01$; ∗∗∗, $p < 0.001$. *n* denotes the number of mice. Detailed information on the test statistics is provided in **Extended Data Table 4**.

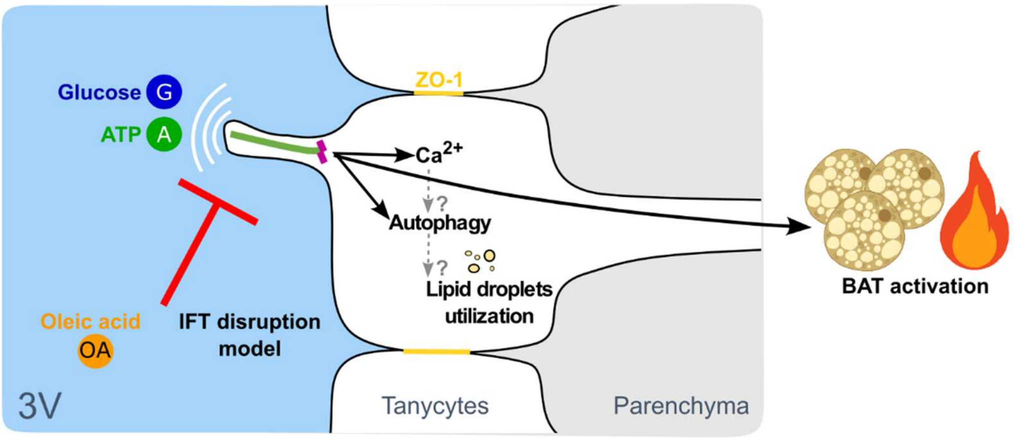

**Extended Data Fig. 5. Tanycytic cilia are key sensory organelles on tanycytes that detect CSF-derived metabolic cues and are essential for maintaining intracellular signalling responses in tanycytes.**

Tanycytic cilia are dynamically regulated by nutrient availability and are essential for proper $Ca^{2+}$ signalling in tanycytes. Excess oleic acid blunts ATP-evoked $Ca^{2+}$ responses and triggers lipid droplet accumulation. Likewise, direct disruption of tanycytic cilia diminishes ATP- and glucose-induced $Ca^{2+}$ signalling, disrupts autophagic flux, and promotes lipid storage. In vivo, tanycytic cilia impairment led to increased body weight and reduced core body temperature and brown adipose tissue activity. Together, these findings identify tanycytic cilia as key regulators of tanycyte metabolic function and underscore their role in whole-body energy homeostasis.

**Extended Data Table 1.**
**Primers used for real-time RT-PCR**

| Gene | Species | Direction | Sequence | Amplificate length [bp] |
|---|---|---|---|---|
| WPRE | Woodchuck hepatitis virus | Fwd | ACTGTGTTTGCTGACGCAAC | 174 |
| | | Rev | CAACACCACGGAATTGTCAG | |
| *Ppia* | Mouse | Fwd | GCATACAGGTCCTGGCATCT | 98 |
| | | Rev | CATCCAGCCATTCAGTCTTGG | |
| *Pgc-1a* | Mouse | Fwd | TCTCAGTAAGGGGCTGGTTG | 151 |
| | | Rev | AGCAGCACACTCTATGTCACTC | |
| *Tfam* | Mouse | Fwd | GAGGCAAAGGATGATTCGGCTC | 116 |
| | | Rev | CGAATCCTATCATCTTTAGCAAGC | |
| *Gk* | Mouse | Fwd | TGGGTAGAACAAGACCCGAAG | 132 |
| | | Rev | TTCCCTCTGGTTGCTGACAC | |
| *Fasn* | Mouse | Fwd | TGCACCTCACAGGCATCAAT | 104 |
| | | Rev | GTCCCACTTGATGTGAGGGG | |
| *Ucp3* | Mouse | Fwd | GAGATGGTGACCTACGACATCA | 158 |
| | | Rev | GCGTTCATGTATCGGGTCTTTA | |
| *Ift88* | Rat | Fwd | CTCAGGCGTCGCGTCTTC | 72 |
| | | Rev | ATCATGTGTACCTCGCGCC | |
| *Kif3a* | Rat | Fwd | TGGAAGGCTACAATGGGACC | 83 |
| | | Rev | CACTGCTCGAACACCTTCCA | |
| *Ppia* | Rat | Fwd | GCATACAGGTCCTGGCATCT | 98 |
| | | Rev | CATCCAGCCATTCAGTCTTGG | |

**Extended Data Table 2.**
Antibodies used for immunofluorescence (IF) and Western blot (WB).

| Antigen | Host species | Source, Catalogue number | Dilution or final concentration | Method |
|---|---|---|---|---|
| ARL13B | Mouse | Antibodies Incorporated / NeuroMab, #73-287 | 1:400<br>1:100 | IF<br>STED |
| Pericentrin | Rabbit | Abcam, #ab4448 | 1:800 | IF |
| AC3 | Rabbit | ThermoFischer Scientific, #PA5-35382 | 1:400<br>1:100 | IF<br>STED |
| IFT88 | Rabbit | Proteintech, #13967-1-AP | 1:400 | IF |
| | | | 1:2000 | WB |
| ZO-1 | Rat | Thermo Fisher Scientific, #14-9776-82 | 1:200 | IF |
| Vimentin | Chicken | Synaptic Systems # 172 006 | 1:500<br>1:400 | IF<br>STED |
| | Chicken | Sigma Aldrich, #AB5733 | 1:500 | IF |
| MAP1LC3B | Rabbit | Sigma-Aldrich, #L7543 | 1:200 | IF |
| | | | 1:20000 | WB |
| p62/SQSTM1 | Guinea pig | Progen, #GP62-N | 1:200 | IF |
| | | | 1:2000 | WB |
| Actin | Mouse | Millipore Sigma, #MAB1501 | 1:2000 | WB |
| Tubulin | Mouse | Sigma-Aldrich, #T9026 | 1:2000 | WB |
| UCP1 | Rabbit | Abcam, #ab10983 | 1:2000 | WB |
| LipiDye™II | / | Funakoshi, #FDV-0027 | 1:1000 | IF |
| Mouse IgG Alexa Fluor 488 | Donkey | ThermoFischer Scientific, #A-21202 | 1:1000 | IF |
| Rabbit IgG Alexa Fluor 555 | Donkey | ThermoFischer Scientific, #A-31572 | 1:1000 | IF |
| Chicken IgG Alexa Fluor 647 | Goat | ThermoFischer Scientific, #A-21449 | 1:1000 | IF |
| Rat IgG Alexa Fluor 647 | Goat | ThermoFischer Scientific, #A-21247 | 1:1000 | IF |
| Rabbit IgG Cy3 | Donkey | Jackson ImmunoResearch Labs, #711-165-152 | 1:1000 | IF |
| Rabbit IgG Alexa Fluor 555 | Goat | ThermoFischer Scientific, #A-21429 | 1:1000 | IF |
| Mouse IgG Alexa Fluor 633 | Goat | ThermoFischer Scientific, #A-21050 | 1:1000 | IF |
| Guinea pig IgG Alexa Fluor 633 | Goat | ThermoFischer Scientific, #A-21105 | 1:1000 | IF |
| Rabbit IgG Alexa Fluor 488 | Donkey | ThermoFischer Scientific, #A-21206 | 1:1000 | IF |
| Mouse IgG | Goat | Abberior, #STORANGE-1001-500UG | 1:800<br>1:400 | IF<br>STED |
| Rabbit IgG | Goat | Abberior, #STRED-1002-500UG | 1:1200<br>1:600 | IF<br>STED |
| Chicken IgG | Goat | Abberior, #STGREEN-1005-500UG | 1:800<br>1:400 | IF<br>STED |

| | | | | |
|---|---|---|---|---|
| Rabbit IgG | Goat | Abberior, #STORANGE-1002-500UG | 1:400 | STED |
| Mouse IgG | Goat | Abberior, #STRED-1001-500UG | 1:600 | STED |
| Mouse IgG HRP | Goat | Jackson ImmunoResearch Labs, #115-035-003 | 1:5000 | WB |
| Rabbit IgG HRP | Goat | DakoCytomation, #P0448 | 1:5000 | WB |
| Guinea pig IgG HRP | Rabbit | ThermoFischer Scientific, #61-4620 | 1:5000 | WB |

**Extended Data Table 3**

**Species and animal characteristics in experiments: genotype, sex, age, viral vectors.**

If not specified otherwise, animals are mice.

| Figure | Species or genotype | Sex | Age | Viral Vectors |
|---|---|---|---|---|
| 1a | C57BL/6 | F | 4m | - |
| 1c | Human | M | 69 years | - |
| 1e, Extended data 1b | C57BL/6 | F | 4m | - |
| 1f, h, l, m<br>Extended data 1e-k | C57BL/6 | M, F | 2m | - |
| 1g, Extended data 1c | C57BL/6 | M, F | 14m | - |
| 1i-k | C57BL/6 | M | 2m 15d<br>16m<br>20m | - |
| Extended data 1l-n | C57BL/6 | F | 2m 17d - 3m 21d | - |
| 2a-l<br>3a-r,<br>Extended data 2a-f | Sprague Dawley rats (primary cultures) | M, F | 10d | - |
| 4a, b, Extended data 3a | *Ift88*$^{fl/fl}$ | M | 4m – 5m | *AAV1/2-Dio2-GFP*<br>*Or*<br>*AAV1/2-Dio2-Cre-2A-GFP* |
| 4c, Extended data 3d | *Ift88*$^{fl/fl}$ | M, F | 4m – 5m | *AAV1/2-Dio2-GFP*<br>*Or*<br>*AAV1/2-Dio2-Cre-2A-GFP* |
| 4d-k, Extended data 3b, Extended data 4a-c | *Ift88*$^{fl/fl}$ | F | 4m – 5m | *AAV1/2-Dio2-GFP*<br>*Or*<br>*AAV1/2-Dio2-Cre-2A-GFP* |
| Extended data 3c, e, f, Extended data 4d-k | *Ift88*$^{fl/fl}$ | M | 4m – 5m | *AAV1/2-Dio2-GFP*<br>*Or*<br>*AAV1/2-Dio2-Cre-2A-GFP* |

**Extended Data Table 4**

**Sample size and results of statistical tests.**

| Figure | Sample size (n) | Statistical test | Values | Comments |
|---|---|---|---|---|
| 1g | n = 4 mice/group | One-way ANOVA followed by Tukey´s post-hoc tests | *F* (3,12) = 41.4, *p* < .001<br><br>α1 vs. α2: *p* = .018<br>α1 vs. β1: *p* < .001<br>α1 vs. β 2: *p* < .001<br>α2 vs. β1: *p* = .027<br>α2 vs. β2: *p* < .001<br>β1 vs. β2: *p* = .012 | |
| 1h | n = 8 mice/group | Welch ANOVA followed by Dunnett´s T3 post-hoc tests | *W* (3,12.1) = 24.4, *p* < .001<br><br>α1 vs. α2: *p* = .119<br>α1 vs. β1: *p* = .004<br>α1 vs. β2: *p* = .002<br>α2 vs. β1: *p* = .016<br>α2 vs. β2: *p* = .006<br>β1 vs. β2: *p* > .999 | |
| 1j | n = mice<br><br>2.5m: 4<br>16m: 5<br>20m: 5 | Scheirer-Ray-Hare test followed by two-tailed Mann-Whitney U post hoc tests, Bonferroni-Holm corrected | age: $\chi^2$ (2) = 6.7, *p* = .005, tanycytes subtype: $\chi^2$ (3) = 42.3, *p* < .001, interaction: $\chi^2$ (6) =.5, *p* = .928<br><br>2.5m vs. 16m:<br>α1: *p* = .048<br>α2: *p* = .064<br>β1: *p* = .189<br>β2: *p* = .048<br><br>2.5m vs. 20m:<br>α1: *p* = .048<br>α2: *p* = .048<br>β1: *p* = .189<br>β2: *p* = .048<br><br>16m vs. 20m:<br>α1: *p* = .151<br>α2: *p* = .841<br>β1: *p* = .69<br>β2: *p* = .31 | |
| 1k | n = mice<br><br>2.5m: 4<br>16m: 5<br>20m: 5 | Two-way ANOVA followed by Sidak's post-hoc tests | age: *F* (2, 44) = 9.8, *p* < .001, tanycytes subtype: *F* (3, 44) = 17.1, *p* < .001, interaction: *F* (6, 44) = 5.3, *p* < .001<br><br>2.5m vs. 16m:<br>α1: *p* < .001<br>α2: *p* = .125<br>β1: *p* > .999<br>β2: *p* = .58<br><br>2.5m vs. 20m:<br>α1: *p* < .001<br>α2: *p* = .134<br>β1: *p* = .247<br>β2: *p* > .999 | |

| | | | | |
|---|---|---|---|---|
| | | | 16m vs. 20m:<br>α1: $p > .999$<br>α2: $p > .999$<br>β1: $p = .223$<br>β2: $p = .267$ | |
| 1m | n = mice<br><br>α1:<br>Males: 8<br>Females: 6<br>α2:<br>Males: 8<br>Females: 6<br>β1:<br>Males: 9<br>Females: 7<br>β2:<br>Males: 9<br>Females: 7 | Scheirer-Ray-Hare test followed by two-tailed Mann-Whitney U post hoc tests, Bonferroni-Holm corrected | sex: $\chi^2$ (1) = 6.8, $p$ = .005, tanycytes subtype: $\chi^2$ (3) = 35.2, $p < .001$, interaction: $\chi^2$ (6) =.5, $p$ = .928<br><br>α1: $p = .414$<br>α2: $p = .06$<br>β1: $p = .06$<br>β2: $p = .031$ | |
| 2b | n = coverslips<br><br>Hypo: 7<br>T3: 7<br>T4: 5<br>TSH: 6 | Kruskal-Wallis followed by Dunn´s post-hoc tests | $\chi^2$ (3) = 9.7, $p$ = .022<br><br>Hypo vs. T3: $p$ = .234,<br>Hypo vs. T4: $p$ = .022,<br>Hypo vs. TSH + PA: $p$ = .03 | 2 independent experiments |
| 2c | n = coverslips<br><br>Hypo: 7<br>T3: 7<br>T4: 5<br>TSH: 6 | One-way ANOVA | $F$ (3,21) = 1.4, $p$ = .273 | 2 independent experiments |
| 2e | n = coverslips<br><br>Growth medium: 6<br>Starvation: 8 | Two-tailed unpaired t-test | $T$ (12) = .1, $p$ = .925 | 3 independent experiments |
| 2f | n = coverslips<br><br>Growth medium: 6<br>Starvation: 8 | Two-tailed unpaired t-test | $T$ (12) = 2.8, $p$ = .015 | 3 independent experiments |
| 2h | n = 4 coverslips / group | Mann-Whitney U tests | Vehicle vs. OA: $p$ = .029,<br>Vehicle vs. PA : $p$ = .886,<br>Vehicle vs. OA + PA: $p$ = .2 | 2 independent experiments |
| 2i | N = 4 coverslips / group | Two-tailed unpaired t-tests | Vehicle vs. OA: $T$ (6) = .7, $p$ = .529,<br>Vehicle vs. PA : $T$ (6) = .3, $p$ = .75,<br>Vehicle vs. OA + PA: $T$ (6) = 2.0, $p$ = .093 | 2 independent experiments |
| 2k | n = coverslips<br><br>Vehicle: 10<br>Oleic acid: 3<br>Palmitic acid: 9 | Kruskal-Wallis followed by Dunn´s post-hoc tests | $\chi^2$ (3) = 23.7, $p$ < .001<br><br>Vehicle vs. OA: $p$ = .034, | One outlier in Palmitic acid group<br><br>2 independent experiments |

| | | | | |
|---|---|---|---|---|
| | Oleic + Palmitic acid: 10 | | Vehicle vs. PA : *p* = .748,<br>Vehicle vs. OA + PA: *p* < .001 | |
| 2l | n = coverslips<br><br>Vehicle: 10<br>Oleic acid: 3<br>Palmitic acid: 10<br>Oleic + Palmitic acid: 10 | Kruskal-Wallis followed by Dunn´s post-hoc tests | $\chi^2$ (3) = 23.2, *p* < .001<br><br>Vehicle vs. OA: *p* = .025,<br>Vehicle vs. PA : *p* > .999,<br>Vehicle vs. OA + PA: *p* < .001 | 2 independent experiments |
| 3c | n = cells<br><br>Vehicle: 350<br>Oleic acid: 299<br>Oleic + Palmitic acid: 300 | Kruskal-Wallis followed by Dunn´s post-hoc tests | $\chi^2$ (2) = 36.55, *p* < .001<br><br>BSA-MeOH vs. OA + PA: *p* = .04,<br>BSA-MeOH vs. OA: *p* < .001,<br>OA + PA vs. OA: *p* = .002 | One outlier in Oleic acid group<br><br>2 independent experiments |
| 3e | n = wells<br><br>Scramble: 4<br>siRNA-*Ift88*: 4 | Two-tailed unpaired t-test | *T* (6) = 3.7, *p* = .01 | |
| 3f | n = wells<br><br>Scramble: 3<br>siRNA-*Kif3a*: 3 | Two-tailed unpaired t-test | *T* (4) = 4.9, *p* = .008 | |
| 3h | n = cells<br><br>Scramble: 146<br>siRNA-*Ift88*: 150<br>siRNA-*Kif3a*: 141 | Kruskal-Wallis followed by Dunn´s post-hoc tests | $\chi^2$ (2) = 35.67, *p* < .001<br><br>Scramble vs. siRNA-Ift88: *p* < .001,<br>Scramble vs. siRNA-Kif3a: *p* < .001,<br>siRNA-Ift88 vs. siRNA-Kif3a: *p* > .999 | 4 outliers in Scramble group, 8 outliers in siRNA-Kif3a group<br><br>3 independent experiments |
| 3j | n = cells<br><br>Scramble: 539<br>siRNA-*Ift88*: 418<br>siRNA-*Kif3a*: 207 | Kruskal-Wallis followed by Dunn´s post-hoc tests | $\chi^2$ (2) = 200, *p* < .001<br><br>Scramble vs. siRNA-Ift88: *p* < .001,<br>Scramble vs. siRNA-Kif3a: *p* < .001,<br>siRNA-Ift88 vs. siRNA-Kif3a: *p* < .001 | 60 outliers in Scramble group, 82 outliers in siRNA-Ift88 group, 43 outliers in siRNA-Kif3a group<br><br>3 independent experiments |
| 3l | n = wells<br><br>Scramble DMSO: 6<br>siRNA-*Ift88* DMSO: 6<br>Scramble BafA1: 6<br>siRNA-*Ift88* BafA1: 6 | Scheirer-Ray-Hare test followed by two-tailed Mann-Whitney U post hoc tests, | Treatment: $\chi^2$ (1) = 15.0, *p* < .001, siRNA: $\chi^2$ (1) = .4, *p* = .527, interaction: $\chi^2$ (1) =2.3, *p* = .134 | 2 independent experiments |

| | | | | |
|---|---|---|---|---|
| | | Bonferroni-Holm corrected | Vehicle + Scramble vs. Bafilomycin+ Scramble: *p* = .009, Vehicle + siRNA-Ift88 vs. Bafilomycin+ siRNA-Ift88: *p* = .03, Vehicle + Scramble vs. Vehicle + siRNA-Ift88: *p* = .026, Bafilomycin + Scramble vs. Bafilomycin + siRNA-Ift88: *p* = .31 | |
| 3m | n = wells<br><br>Scramble DMSO: 6<br>siRNA-*Ift88* DMSO: 6<br>Scramble BafA1: 5<br>siRNA-*Ift88* BafA1: 6 | Two-way ANOVA followed by Sidak's post-hoc tests | treatment: *F* (1, 19) = .7, *p* = .423, siRNA: *F* (1, 19) = 13.0, *p* = .002, interaction: *F* (1, 19) = .5, *p* = .493<br><br>Scramble vs. siRNA-Ift88:<br>Vehicle: *p* = .011<br>Bafilomycin: *p* = .114 | 1 outlier in Scramble – Bafilomycin group<br><br>2 independent experiments |
| 3n | n = wells<br><br>Scramble: 6<br>siRNA-*Ift88*: 6 | Two-tailed unpaired t-test | *T* (10) = 2.3, *p* = .047 | |
| 3p | n = fields of view in 3 coverslips<br><br>Scramble: 18<br>siRNA-*Ift88*: 15 | Mann-Whitney U test | *U* = 30, *p* < .001 | |
| 3q | n = fields of view in 3 coverslips<br><br>Scramble: 19<br>siRNA-*Ift88*: 15 | Mann-Whitney U test | *U* = 66, *p* = .007 | |
| 3h | n = fields of view in 3 coverslips<br><br>Scramble: 19<br>siRNA-*Ift88*: 15 | Mann-Whitney U test | *U* = 80, *p* = .029 | |
| 4c | n = mice<br><br>$Ift88^{fl/fl}$: 22<br>$Ift88^{TanKO}$: 21 | Two-tailed unpaired t-test | *T* (41) = 3.7, *p* < .001 | |
| 4d | n = mice<br><br>$Ift88^{fl/fl}$: 9<br>$Ift88^{TanKO}$: 8 | Two-way repeated measures ANOVA followed by Sidak's post-hoc tests | week: *F* (2.9, 43.1) = 7.8, *p* < .001, AAV: *F* (1, 15) = 5.5, *p* = .034, interaction: *F* (6, 90) = 1.6, *p* = .147 | |
| 4e | n = mice<br><br>$Ift88^{fl/fl}$: 8<br>$Ift88^{TanKO}$: 8 | Mann-Whitney U test | *U* = 4.5, *p* = .002 | 1 outlier in $Ift88^{fl/fl}$ group |
| 4f left | n = mice<br><br>$Ift88^{fl/fl}$: 8<br>$Ift88^{TanKO}$: 8 | Two-way repeated measures ANOVA followed | time: *F* (1.8, 25.6) = 54.3, *p* < .001, AAV: *F* (1, 14) = 3.3, *p* = .089, interaction: *F* (3, 42) = 1.8, *p* = .171 | |

| | | | |
|---|---|---|---|
| | | by Sidak's post-hoc tests | 15 min: *p* = .788<br>30 min: = .289<br>60 min: *p* = .173<br>120 min: *p* > .999 |
| 4f right | n = mice<br><br>*Ift88*$^{fl/fl}$: 8<br>*Ift88*$^{TanKO}$: 8 | Two-tailed unpaired t-test | *T* (14) = 1.8, *p* = .088 |
| 4h | n = mice<br><br>*Ift88*$^{fl/fl}$: 9<br>*Ift88*$^{TanKO}$: 8 | Two-tailed unpaired t-test | *T* (15) = 2.3, *p* = .037 |
| 4i | n = mice<br><br>*Ift88*$^{fl/fl}$: 9<br>*Ift88*$^{TanKO}$: 8 | Mann-Whitney U test | *U* = 23.5, *p* = .246 |
| 4j<br><br>*Pgc1α*<br><br><br>*Tfam*<br><br><br>*GK*<br><br><br>*Fasn*<br><br><br>*UCP3* | n = mice<br><br>*Ift88*$^{fl/fl}$: 8<br>*Ift88*$^{TanKO}$: 7<br><br>*Ift88*$^{fl/fl}$: 8<br>*Ift88*$^{TanKO}$: 7<br><br>*Ift88*$^{fl/fl}$: 8<br>*Ift88*$^{TanKO}$: 7<br><br>*Ift88*$^{fl/fl}$: 8<br>*Ift88*$^{TanKO}$: 7<br><br>*Ift88*$^{fl/fl}$: 5<br>*Ift88*$^{TanKO}$: 4 | Two-tailed unpaired t-tests | Pgc1α: *T* (13) = 1.9, *p* = .079,<br><br>Tfam : *T* (13) = 3.7, *p* = .003,<br><br>GK: *T* (13) = 3.1, *p* = .009,<br><br>Fasn: *T* (13) = 2.2, *p* = .045,<br><br>UCP3: *T* (7) = 2.1, *p* = .077 |
| 4k | n = mice<br><br>*Ift88*$^{fl/fl}$: 9<br>*Ift88*$^{TanKO}$: 8 | Two-tailed unpaired t-test | *T* (15) = 2.2, *p* = .049 |
| Extended data 1c | n = 4 mice/group | One-way repeated measures ANOVA | *F* (1.2,3.4) = 2.9, *p* = .178 |
| Extended data 1e | n = mice<br><br>Males: 4<br>Females: 5 | Two-way repeated measures ANOVA | region: *F* (2, 14) = 35.3, *p* < .001, sex: *F* (1, 7) = .1, *p* = .786, interaction: *F* (2, 14) = 2.4, *p* = .131 |
| Extended data 1f | n = mice<br><br>Males: 5<br>Females: 4 | Two-way repeated measures ANOVA followed by Sidak's post-hoc tests | region: *F* (3, 21) = 106.1, *p* < .001, sex: *F* (1, 7) = 6.2, *p* = .041, interaction: *F* (3, 21) = 1.4, *p* = .264<br><br>α1: *p* = .208<br>α2: *p* = .031<br>β1: *p* = .287<br>β2: *p* = .968 |
| Extended data 1g | n = mice<br><br>Males: 2-4 | - | - |

| | | | |
|---|---|---|---|
| | Females: 1-2 | | |
| Extended data 1h | n = mice<br><br>Males: 4<br>Females: 5 | Two-way repeated measures ANOVA | region: $F$ (2, 14) = .4, $p$ = .71, sex: $F$ (1, 7) = 1.2, $p$ = .302, interaction: $F$ (2, 14) = 1.5, $p$ = .255 |
| Extended data 1i | n = mice<br><br>Males: 5<br>Females: 4 | Scheirer-Ray-Hare test | sex: $\chi^2$ (1) = 1.7, $p$ = .19, region: $\chi^2$ (3) = 1.4, $p$ = .706, interaction: $\chi^2$ (3) =*.1*, $p$ = .996 |
| Extended data 1j | n = mice<br><br>Males: 5<br>Females: 4 | Two-way repeated measures ANOVA | region: $F$ (3, 21) = 28.9, $p$ < .001, sex: $F$ (1, 7) = .6, $p$ = .455, interaction: $F$ (3, 21) = 1.7, $p$ = .197 |
| Extended data 1k | n = mice<br><br>Males: 2-4<br>Females: 1-2 | - | - |
| Extended data 1m | n = mice<br><br>Proestrus: 4<br>Estrus: 3<br>Diestrus: 3 | Two-way repeated measures ANOVA followed by Tukey´s post-hoc tests | region: $F$ (2.6, 17.9) = 82.5, $p$ < .001, cycle phase: $F$ (2, 7) = 11.0, $p$ = .007, interaction: $F$ (6, 21) = 1.3, $p$ = .308<br><br>α1:<br>proestrus vs. estrus: $p$ = .659,<br>proestrus vs. diestrus: $p$ = .063,<br>estrus vs. diestrus: $p$ = .22<br><br>α2:<br>proestrus vs. estrus: $p$ = .343,<br>proestrus vs. diestrus: $p$ = .118,<br>estrus vs. diestrus: $p$ = .036<br><br>β1:<br>proestrus vs. estrus: $p$ = .54,<br>proestrus vs. diestrus: $p$ = .251,<br>estrus vs. diestrus: $p$ = .142<br><br>β2:<br>proestrus vs. estrus: $p$ = .933,<br>proestrus vs. diestrus: $p$ = .807,<br>estrus vs. diestrus: $p$ = .819 |
| Extended data 1n | n = mice<br><br>Proestrus: 4<br>Estrus: 3<br>Diestrus: 3 | RM Scheirer-Ray-Hare test followed by two-tailed Mann-Whitney U post | region: $\chi^2$ (3) = 9.1, $p$ = .028, cycle phase: $\chi^2$ (2) = 3.8, $p$ = .153, interaction: $\chi^2$ (6) = 8.5, $p$ = .206 |

| | | | | |
|---|---|---|---|---|
| | | hoc tests, Bonferroni-Holm corrected | α1:<br>proestrus vs. estrus: *p* > .999,<br>proestrus vs. diestrus: *p* = .686,<br>estrus vs. diestrus: *p* > .999<br><br>α2:<br>proestrus vs. estrus: *p* > .999,<br>proestrus vs. diestrus: *p* = .686,<br>estrus vs. diestrus: *p* > .999<br><br>β1:<br>proestrus vs. estrus: *p* > .999,<br>proestrus vs. diestrus: *p* > .999,<br>estrus vs. diestrus: *p* > .999<br><br>β2:<br>proestrus vs. estrus: *p* > .999,<br>proestrus vs. diestrus: *p* > .999,<br>estrus vs. diestrus: *p* > .999 | |
| Extended data 2a | n = wells<br><br>Scramble: 3<br>siRNA-*Ift88*: 3 | Two-tailed unpaired t-test (Welch corrected) | *T* (2.0) = 6.2, *p* = .025 | |
| Extended data 2b | n = wells<br><br>Scramble: 4<br>siRNA-*Kif3a*: 4 | Two-tailed unpaired t-test | *T* (6) = 4.0, *p* = .007 | |
| Extended data 2c | n = wells<br><br>Scramble: 9<br>siRNA-*Ift88*: 9<br>siRNA-*Kif3a*: 8 | One-way ANOVA followed by Dunnett´s post-hoc tests | *F* (2,23) = 3.2, *p* = .059,<br><br>Scramble vs. siRNA-Ift88: *p* = .716,<br>Scramble vs. siRNA-Kif3a: *p* = .039 | 3 independent experiments |
| Extended data 2e | n = fields of view in 3 coverslips<br><br>Scramble DMSO: 15<br>siRNA-*Ift88* DMSO: 15<br>Scramble BafA1: 15<br>siRNA-*Ift88* BafA1: 12 | Scheirer-Ray-Hare test followed by two-tailed Mann-Whitney U post hoc tests, Bonferroni-Holm corrected | Treatment: $\chi^2$ (1) = 7.9, *p* = .005, siRNA: $\chi^2$ (1) = 10.3, *p* = .001, interaction: $\chi^2$ (1) =.2, *p* = .663<br><br>Vehicle + Scramble vs. Bafilomycin+ Scramble: *p* = .015,<br>Vehicle + siRNA-Ift88 vs. Bafilomycin+ siRNA-Ift88: *p* = .339,<br>Vehicle + Scramble vs. Vehicle + siRNA-Ift88: *p* = .042, | |

| | | | | |
|---|---|---|---|---|
| | | | Bafilomycin + Scramble vs. Bafilomycin + siRNA-Ift88: $p$ = .015 | |
| Extended data 2f | n = fields of view in 3 coverslips<br><br>Scramble DMSO: 15<br>siRNA-*Ift88* DMSO: 14<br>Scramble BafA1: 13<br>siRNA-*Ift88* BafA1: 11 | Scheirer-Ray-Hare test followed by two-tailed Mann-Whitney U post hoc tests, Bonferroni-Holm corrected | Treatment: $\chi^2$ (1) = .3, $p$ = .566, siRNA: $\chi^2$ (1) = 8.4, $p$ = .004, interaction: $\chi^2$ (1) =.6, $p$ = .431<br><br>Vehicle + Scramble vs. Bafilomycin+ Scramble: $p$ = .317, Vehicle + siRNA-Ift88 vs. Bafilomycin+ siRNA-Ift88: $p$ = .65, Vehicle + Scramble vs. Vehicle + siRNA-Ift88: $p$ = .147, Bafilomycin + Scramble vs. Bafilomycin + siRNA-Ift88: $p$ = .285 | 1 outlier<br><br>2 outliers<br>1 outlier |
| Extended data 3b | n = mice<br><br>*Ift88*$^{fl/fl}$: 9<br>*Ift88*$^{TanKO}$: 9 | Two-tailed unpaired t-test | $T$ (16) = 2.3, $p$ = .033 | 1 outlier<br>1 outlier |
| Extended data 3c | n = mice<br><br>*Ift88*$^{fl/fl}$: 13<br>*Ift88*$^{TanKO}$: 12 | Two-tailed unpaired t-test | $T$ (23) = 2.7, $p$ = .012 | 1 outlier<br>1 outlier |
| Extended data 3d | n = mice<br><br>*Ift88*$^{fl/fl}$: 7<br>*Ift88*$^{TanKO}$: 7 | Two-tailed unpaired t-test | $T$ (12) = 2.3, $p$ = .04 | |
| Extended data 3e | n = mice<br><br>*Ift88*$^{fl/fl}$: 3<br>*Ift88*$^{TanKO}$: 3 | Two-way repeated measures ANOVA | tanycyte subtype: $F$ (3, 12) = 55.0, $p$ < .001, rAAV: $F$ (1, 4) = .04, $p$ = .853, interaction: $F$ (3, 12) = 2.2, $p$ = .138 | |
| Extended data 3f | n = mice<br><br>*Ift88*$^{fl/fl}$: 3<br>*Ift88*$^{TanKO}$: 3 | Two-way repeated measures ANOVA | tanycyte subtype: $F$ (3, 12) = 1.5, $p$ = .261, rAAV: $F$ (1, 4) = .4, $p$ = .56, interaction: $F$ (3, 12) = .5, $p$ = .724 | |
| Extended data 4a | n = mice<br><br>*Ift88*$^{fl/fl}$: 9<br>*Ift88*$^{TanKO}$: 8 | Two-way repeated measures ANOVA | time: $F$ (1.2, 18.4) = 178.6, $p$ < .001, rAAV: $F$ (1, 15) = .2, $p$ = .709, interaction: $F$ (23, 345) = .4, $p$ = .991 | |
| Extended data 4b | n = mice<br><br>*Ift88*$^{fl/fl}$: 9<br>*Ift88*$^{TanKO}$: 8 | Two-way repeated measures ANOVA | time: $F$ (1.6, 24.6) = 9.8, $p$ = .001, rAAV: $F$ (1, 15) = 2.4, $p$ = .143, interaction: $F$ (3, 45) = .1, $p$ = .946 | |

| | | | | |
|---|---|---|---|---|
| Extended data 4c | n = mice<br><br>*Ift88$^{fl/fl}$*: 9<br>*Ift88$^{TanKO}$*: 8 | Two-way repeated measures ANOVA | time: $F$ (1.8, 26.2) = 35.5, $p$ < .001, rAAV: $F$ (1, 15) = 3.8, $p$ = .071, interaction: $F$ (3, 45) = .3, $p$ = .818 | |
| Extended data 4d | n = mice<br><br>*Ift88$^{fl/fl}$*: 10<br>*Ift88$^{TanKO}$*: 11 | Two-way repeated measures ANOVA | time: $F$ (3.2, 60.6) = 11.2, $p$ < .001, rAAV: $F$ (1, 19) = .1, $p$ = .735, interaction: $F$ (6, 114) = .1, $p$ = .996 | 1 outlier |
| Extended data 4e | n = mice<br><br>*Ift88$^{fl/fl}$*: 10<br>*Ift88$^{TanKO}$*: 11 | Mann-Whitney U test | $U$ = 30.5, $p$ = .087 | 1 outlier |
| Extended data 4f left | n = mice<br><br>*Ift88$^{fl/fl}$*: 11<br>*Ift88$^{TanKO}$*: 9 | Two-way repeated measures ANOVA | time: $F$ (1.6, 28.8) = 78.6, $p$ < .001, rAAV: $F$ (1, 18) = .8, $p$ = .387, interaction: $F$ (3, 54) = .1, $p$ = .96 | |
| Extended data 4f right | n = mice<br><br>*Ift88$^{fl/fl}$*: 11<br>*Ift88$^{TanKO}$*: 9 | Two-tailed unpaired t-test | $T$ (18) = .5, $p$ = .64 | |
| Extended data 4g | n = mice<br><br>*Ift88$^{fl/fl}$*: 11<br>*Ift88$^{TanKO}$*: 10 | Two-way repeated measures ANOVA | time: $F$ (1.2, 22.9) = 251.3, $p$ < .001, rAAV: $F$ (1, 19) = .9, $p$ = .931, interaction: $F$ (23, 437) = .2, $p$ > .999 | |
| Extended data 4h | n = mice<br><br>*Ift88$^{fl/fl}$*: 11<br>*Ift88$^{TanKO}$*: 10 | Two-way repeated measures ANOVA | time: $F$ (1.3, 24.5) = 117.4, $p$ < .001, rAAV: $F$ (1, 19) = .3, $p$ = .603, interaction: $F$ (3, 57) = 1.3, $p$ = .292 | |
| Extended data 4i | n = mice<br><br>*Ift88$^{fl/fl}$*: 11<br>*Ift88$^{TanKO}$*: 10 | Two-way repeated measures ANOVA | time: $F$ (1.9, 36.7) = 228.4, $p$ < .001, rAAV: $F$ (1, 19) = .02, $p$ = .888, interaction: $F$ (3, 57) = .9, $p$ = .442 | |
| Extended data 4j | n = mice<br><br>*Ift88$^{fl/fl}$*: 11<br>*Ift88$^{TanKO}$*: 11 | Two-tailed unpaired t-test | $T$ (20) = .6, $p$ = .525 | |
| Extended data 4k | n = mice<br><br>*Ift88$^{fl/fl}$*: 11<br>*Ift88$^{TanKO}$*: 11 | Mann-Whitney U test | $U$ = 48, $p$ = .439 | |
